# Assignment of structural transitions during mechanical unwrapping of nucleosomes and their disassembly products

**DOI:** 10.1101/2022.04.13.488252

**Authors:** César Díaz-Celis, Cristhian Cañari-Chumpitaz, Robert P. Sosa, Juan P. Castillo, Meng Zhang, Enze Cheng, Andy Chen, Michael Vien, JeongHoon Kim, Bibiana Onoa, Carlos Bustamante

## Abstract

Nucleosome DNA unwrapping and its disassembly into hexasomes and tetrasomes is necessary for genomic access and plays an important role in transcription regulation. Previous single-molecule mechanical nucleosome unwrapping revealed a low- and a high-force transitions, and force-FRET pulling experiments showed that DNA unwrapping is asymmetrical occurring always first from one side before the other. However, the assignment of DNA segments involved in these transitions remains controversial. Here, using high-resolution optical tweezers with simultaneous single-molecule FRET detection we show that the low-force transition corresponds to the undoing of the outer-wrap of one side of the nucleosome (~27 bp), a process that can occur either cooperatively or non-cooperatively, whereas the high-force transition corresponds to the simultaneous unwrapping of ~76 bp from both sides. This process may give rise stochastically to the disassembly of nucleosomes into hexasomes and tetrasomes whose unwrapping/rewrapping trajectories we establish. In contrast, nucleosome rewrapping does not exhibit asymmetry. To rationalize all previous nucleosome unwrapping experiments, it is necessary to invoke that mechanical unwrapping involves two nucleosome reorientations: one that contributes to the change in extension at the low-force transition, and another that coincides but does not contribute to the high-force transition.

**Significance statement:** Nucleosomes, the fundamental structural unit of chromatin, consists of ~147 DNA base pairs wrapped around a histone protein octamer. Determining the forces required to unwrap the DNA from the core particle and the stepwise transitions involved in the process are essential to characterize the strength of the nucleosomal barrier and its contribution as a mechanism of control of gene expression. Here, we performed combined optical tweezers and single-molecule fluorescence measurements to annotate the specific DNA segments unwrapping during the force transitions observed in mechanical unwrapping of nucleosomes. Furthermore, we characterize the mechanical signatures of subnucleosomal particles: hexasomes and tetrasomes. The characterization performed in this work is essential for the interpretation of ongoing studies of chromatin remodelers, polymerases, and histone chaperones.

## Introduction

Chromatin is a nucleoprotein complex that regulates DNA accessibility for replication, repair, and transcription in eukaryotic cells. The structural unit of chromatin is the nucleosome consisting of 147 bp DNA wrapped in 1.65 left-handed turns around an octamer of histone proteins (1, 2). The octamer core is composed of two copies of histones H2A, H2B, H3, and H4, organize in two H2A-H2B heterodimers and in one (H3-H4)_2_ tetramer. The histone core is stabilized mainly by hydrophobic attractions, whereas DNA-histone contacts involve hydrogen bonds and electrostatic interactions (2). Nucleosomes are highly dynamic and intrinsically plastic complexes which are subject to extensive modifications to regulate access to DNA, including exchange of canonical histones with histone variants, histone post-translational modifications, and interactions with regulatory proteins such as histone chaperones and molecular motors (RNA polymerase, remodelers, etc.). These modifications alter histone-DNA interactions leading to nucleosome disassembly into subnucleosomal particles—hexasomes and tetrasomes—and the exposure of DNA (3–8)). Accordingly, a detailed picture of nucleosomal regulation of gene expression requires a precise characterization of the mechanical stability of the nucleosome and how fluctuations, forces, and torques affect its integrity.

The binding of DNA to the histone core and the structural transitions that occur during nucleosome disassembly have traditionally been studied by bulk approaches (9–13). An alternative approach is the use of single-molecule mechanical manipulation with optical or magnetic trapping. These methods provide unique insight into the process by which the cellular machinery access the nucleosome-bound DNA and the effect of epigenetics marks on nucleosome accessibility. These experiments include the mechanical stretching of chromatin fibers and nucleosome arrays (14–20), and of single nucleosomes (21–24). Likewise, torque has been applied to nucleosomes using magnetic trapping (25, 26) and torsional optical tweezers (27).

These single-molecule force-extension experiments revealed that under applied force, DNA unwraps from the histone core in two stages. The first, occurring between 3 and 6 pN, was assigned to the unwrapping of the outer DNA turn releasing 60-70 bp in a reversible manner. The second, taking place over a broad range of forces (8-40 pN), is characterized by an abrupt change in extension or rip, and was attributed to the unwrapping of the DNA inner turn (~80 bp) (16, 17, 21, 23, 24, 27). Even though the estimated amount of unwrapped DNA described in these studies was in agreement with the nucleosome crystal structure, it assumed that unwrapping under tension occurs symmetrically from both ends.

In contrast, combined single-molecule fluorescence-force spectroscopy experiments of FRET pair-labeled nucleosomes revealed that the low-force transition corresponds only to the unwrapping of one outer wrap DNA arm, whereas the high-force transition corresponded to the unidirectional unwrapping of internal nucleosome positions and the other arm. Thus, according to these studies, DNA unwrapping under tension is asymmetrical and occurs from the weak towards the strong arm (22). Although these single-molecule fluorescence experiments targeted specific regions of nucleosomal DNA, they were not able to directly measure the changes in extension associated with each force transition due to limited resolution. Taken together, the interpretation of the changes in extension observed by previous pulling experiments remain inconclusive. Likewise, the mechanical signatures of hexasomes and tetrasomes have not been established.

Here we use high-resolution optical tweezers with single-molecule fluorescence detection capability (“fleezers”) to characterize the unwrapping and rewrapping of nucleosomes, hexasomes, and tetrasomes under tension. We assign each force transition to the unwrapping of specific DNA regions and monitor the integrity of nucleosomes during their mechanical unwrapping, finding that their disassembly is stochastic. Our results reconcile previous discrepancies in the interpretation of single-molecule mechanical unwrapping of nucleosomes.

## Results

### Mechanical signatures of nucleosome unwrapping

We assembled nucleosomes by salt dialysis using a recombinant *Xenopus laevis* histone core and the artificial 601 nucleosome-positioning sequence (601 NPS), and purified them using polyacrylamide gel electrophoresis (Fig. 1A). To perform the mechanical experiments, nucleosomes were ligated to DNA-handles and anchored to 1 µm polystyrene microbeads that were trapped in the high-resolution optical tweezers instrument (Fig. 1B).

**Fig. 1.**
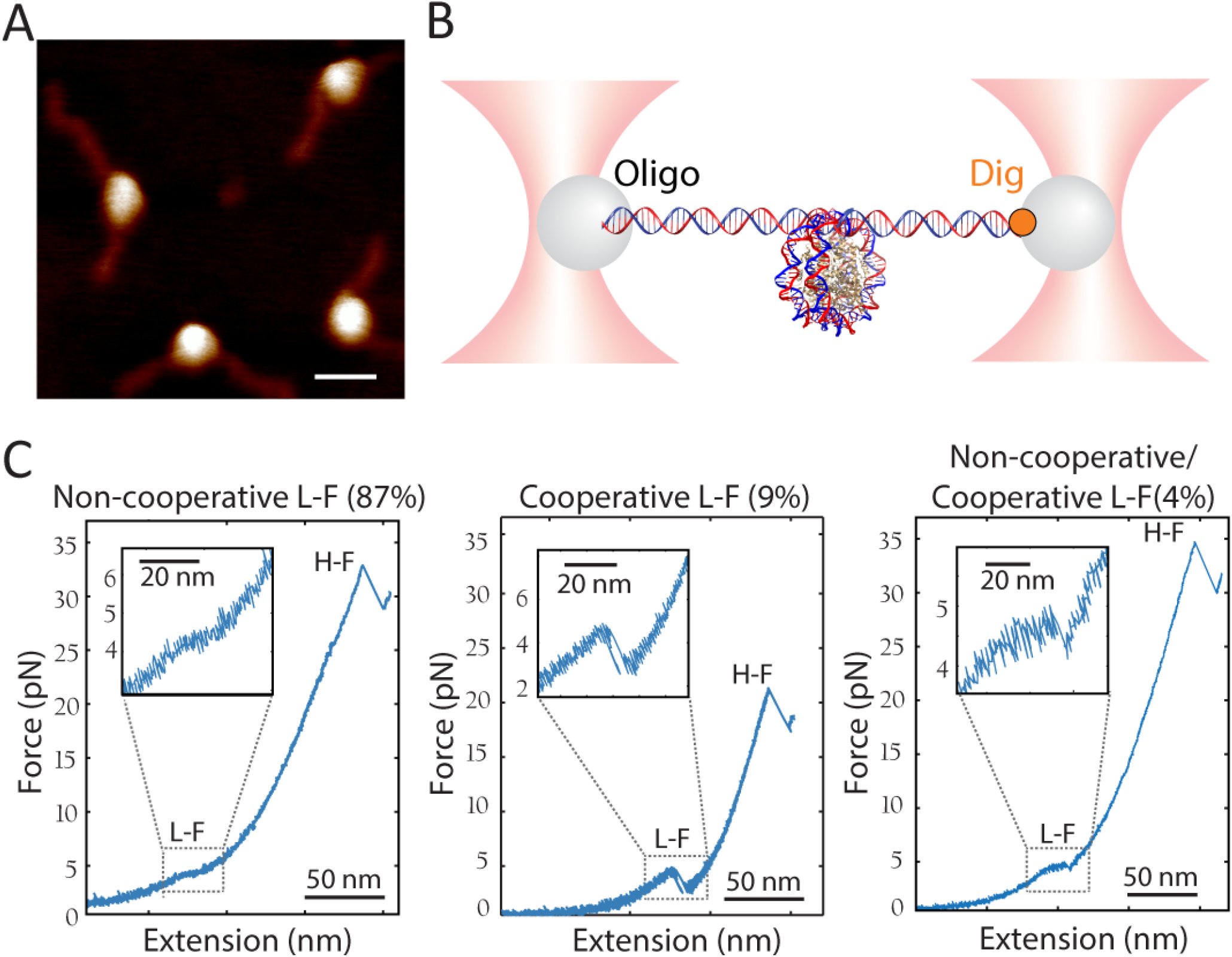
Nucleosome unwrapping trajectory under tension. (A) Atomic force microscopy of purified *X. laevis* nucleosomes reconstituted on 601 NPS and flanked by 100 bp and 50 bp of DNA. Nucleosome average maximum height corresponded to ~3.9 nm (n= 4). Scale bar= 20 nm. (B) Experimental geometry for single nucleosome manipulation using high-resolution optical tweezers. Nucleosome was ligated to DNA handles, which in turn were tethered to polystyrene microbeads by DNA ligation with DNA oligo coated beads, and by biotin binding to streptavidin coated beads. (C) Examples of force-extension unwrapping trajectories of single nucleosomes. L-F denotes the low-force transition. H-F denotes the high-force transition. The low-force transition can be non-cooperative (left panel), cooperative (middle panel), or exhibit a combination of both mechanisms of unwrapping (right panel).

At low ionic strength (50 mM KOAc), pulling curves showed the two force-extension transitions previously described for single nucleosomes at low- and at high-force (21, 23) (Fig. 1C). In our conditions the low-force transition (N= 53) manifests either as: i) a non-cooperative transition at 3.9 ± 0.3 pN, observed in the majority of the pulling curves (88%; n= 45) and that appears as a shoulder or plateau in the force-extension curve with an extension of 19 ± 1 nm (Fig. 1C; left panel); ii) a cooperative transition at 4.2 ± 0.8 pN (8%; n= 4) characterized by an abrupt change in extension or rip of 20.4 ± 1.5 nm (Fig. 1C; middle panel); and iii) a combination of the cooperative and non-cooperative transitions at 3.8 ± 0.2 pN (4%; n= 2) with a change in extension of 20.6 ± 2.0 nm (Fig. 1C; right panel).

In previous studies using reconstituted chicken erythrocyte nucleosomes, the low-force transition at 50 mM KOAc (21) and at 10 mM NaN_3_ (23) appeared as a rip and became progressively less cooperative as the ionic strength was increased to 200 mM KOAc (21). To determine if the ionic strength modifies the unwrapping trajectory of the low-force transition, we stretched nucleosomes at a lower (~10 mM K^+^ from KOH) and at a higher (200 mM KOAc) concentrations (Fig. S1A and B; Table S1). We observed no change in the frequency (□ 85%) nor the extension of the non-cooperative transition with the increased ionic strength, while the force decreases slightly (Table S1).

Because nucleosome stability is depends on the DNA sequence, we analyzed the low-force transition of *X. laevis* nucleosomes assembled on the natural 5S rRNA NPS, which unlike the artificial 601 NPS (28, 29), has a lower affinity for the octamer, resulting in less accurate positioning and stability (30). In 50 mM KOAc, 98% of the low-force transitions of the 5S nucleosomes are non-cooperative (Fig. S1C; Table S1), indicating that the NPS has a small effect on the low-force unwrapping trajectories. Likewise, the low-force transition is invariant to the recombinant nucleosome source (*X. laevis*, human (Fig. S2A), or yeast (Fig. S2B) and Table S1).

The second force-extension transition of nucleosome unwrapping is characterized by a rip that occurs abruptly over a broad range of forces above ~12 pN up to ~ 37 pN (Fig. 1C; Table S2). At 50 mM KOAc, the high-force transition occurs at 30.4 ± 8.3 pN—higher than the corresponding transition observed with nucleosomes assembled with native histones (21)—and exhibits a change in extension of 24.5 ± 1.5 nm. The ionic strength does not affect the cooperativity of this transition nor the change in extension (Table S2); at 10 mM K^+^ from KOH, the high-force rip is centered at ~35 pN (Fig. S1A; Table S2), while at 200 mM KOAc the high-force rip occurs at ~24 pN (Fig. S1B, Table S2). The reduction of the unwrapping force with ionic strength agrees with the electrostatic nature of DNA-histone interactions. In the case of *X. laevis* 5S (Fig. S1C) and human 601 nucleosomes (Fig. S2A), the high-force transition at 50 mM KOAc is seen at ~29 pN, while for yeast 601 nucleosomes it occurs at ~21 pN (Fig. S2B). These observations suggest different histone-DNA interactions between recombinant nucleosomes from vertebrates and invertebrates. Despite these differences in force, the change in extension is similar for all three types of recombinant nucleosomes (Table S2).

To summarize, the majority of trajectories of recombinant nucleosomes displayed a non-cooperative low-force transition and a high-force rip centered at high forces, in contrast to those of nucleosomes assembled with native histones which are characterized by a cooperative low-force transition and a high-force rip below ~15 pN (21, 31). These discrepancies may be due to the presence of post-translational modifications present in the native octamers, some of which have been shown to affect the mechanical unwrapping of nucleosome arrays (16, 19).

### Mechanical unwrapping o nucleosomes generates hexasomes and tetrasomes

*In vitro* studies have shown that nucleosome destabilization at increasing concentration of salt proceeds via DNA unwrapping and the sequential dissociation of H2A-H2B heterodimers, yielding hexasomes and tetrasomes (9–13, 32). However, these intermediates have not been identified in mechanical disassembly experiments, nor their unwrapping trajectories characterized.

To investigate the disassembly of nucleosomes under force, we subjected single nucleosomes to successive pulling/relaxation cycles, from low-to high-force and back, until we obtained the force-extension curve of bare DNA. The first pulling curve in Fig. 2A, represents the intact nucleosome and shows the two force-extension transitions previously described. The second and third cycles display two distinct types of unwrapping trajectories with a single force-extension signature each. The first type of trajectory (Fig. 2A, “type I”) exhibits a rip or cooperative transition centered at 23.8 ± 4.3 pN (n= 42) with a change in extension of 23.4 ± 1.1 nm. The second or “type II” trajectory typically precedes the bare DNA force-extension signature and exhibits a transition at low-force (4.5 ± 0.8 pN; n= 40), a net change in extension of 14.9 ± 1.7 nm and a plateau-shape that displayed dynamic fluctuations (hopping). Disassembly of *X. laevis* 5S nucleosomes, as well as human and yeast 601 nucleosomes, also produces type I and type II pulling trajectories (Fig. S3A-C).

**Fig 2.**
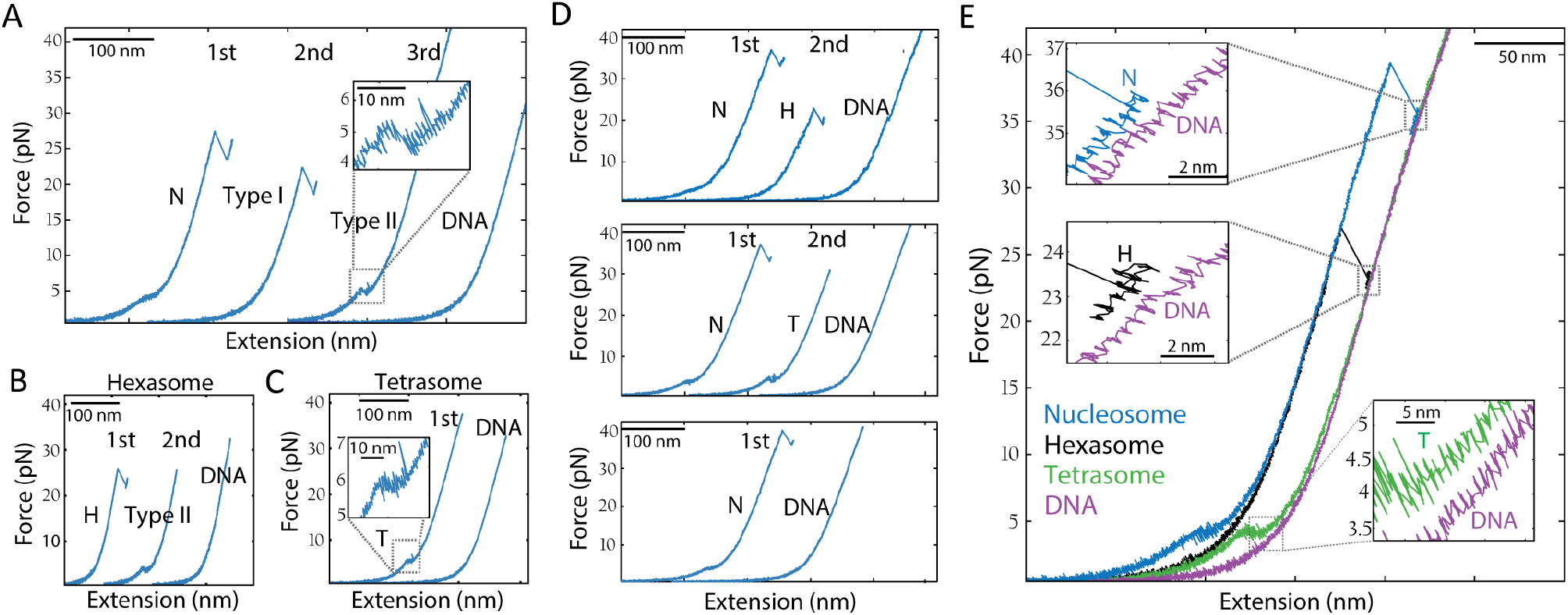
Nucleosome disassembly under force generates hexasomes and tetrasomes and it is stochastic. (A) Nucleosome, (B) hexasome, and (C) tetrasome disassembly by pulling and relaxation cycles (1^st^, 2^nd^, and 3rd) at 50 mM KOAc. Only pulling curves (blue) are shown and they were artificially separated for illustrative purposes. (A) The first pulling curve from the first cycle correspond to the nucleosome (N). Type I and type II intermediates were observed in the pulling curves of the second and third cycles, respectively. Type I exhibit a single rip at ~22 pN and type II exhibit a transition with fluctuations or hopping at ~4.5 pN (inset). After third cycle was completed, the next pulling curve corresponded to bare DNA, indicating full nucleosome disassembly. (B) Hexasome (H) unwrapping trajectory exhibit a single force-extension transition as a rip (~25 pN). Type II intermediate is generated in the second cycle before full disassembly (DNA pulling curve). (C) Tetrasome (T) unwrapping trajectory exhibit a single force-extension transition with hopping (inset) at a low-force of ~5 pN. (D) Mechanical disassembly of nucleosomes is stochastic. Under force, nucleosomes (N) can disassembly into: an hexasome without forming tetrasomes before full dissociation (left panel); into a tetrasome (middle panel); or fully dissociate in one pulling and relaxation cycle (right panel). (E) Pulling trajectories of nucleosomes, hexasomes, tetrasomes, and DNA.

To establish the identity of the type I and type II pulling trajectories, we purified hexasomes and tetrasomes and characterized them by atomic-force microscopy (Fig. S4A and B). The first pulling curve of hexasomes (Fig. 2B) showed an unwrapping trajectory indistinguishable from the type I trajectory (Fig. 2A), displaying a single rip of 23.4 ± 1.1 nm at a force of 24.6 ± 3.0 pN. Successive pulling/relaxation cycles of hexasomes also generate type II and bare DNA trajectories, in that order. Force-extension trajectories of tetrasomes (Fig. 2C) exhibit a single transition of 13.3 ± 1.7 nm at a force of 4.4 ± 0.5 pN, resembling the type II trajectories. Interestingly, pulling trajectories of nucleosomes, hexasomes, and tetrasomes were not always observed consecutively through pulling/relaxation cycles (Fig. 2A): a nucleosome can also disassemble directly into tetrasomes or bare DNA with different probabilities depending on the ionic strength (Fig. 2D; Table S3), indicating that the mechanical disassembly is stochastic. Fig. 2E depicts the characteristic force-extension curves of nucleosomes, hexasomes, tetrasomes and bare DNA. Notice that after the last unwrapping transition, the pulling curve of bare DNA is longer by 2.3 ± 0.6 nm (n= 20), 2.5 ± 0.8 nm (n= 14), and 7.8 ± 1.7 nm (n= 17), than those of the nucleosome, hexasome, and tetrasome, respectively, indicating that even at forces > 30 pN part of the DNA remains wrapped around the histone core.

Previous stretching experiments using torsional optical tweezers of tetrasomes assembled with (H3-H4)_2_ tetramers reported pulling curves displaying a single rip at ~16 pN (27), that resemble those of hexasomes described here (Fig. 2B), and quite different from the pulling curves of tetrasomes characterized also in this study (Fig. 2C). We proposed that this discrepancy results from (H3-H4)_2_ tetramer oligomerization into nucleosome-like particles as previously described (33–35). Indeed, using AFM, we found that whereas at 1:1.4 (H3-H4)_2_:DNA ratio, (H3-H4)_2_ tetramers form mainly tetrasomes, at higher (H3-H4)_2_:DNA ratios (1:2.6), they assemble into bigger particles resembling hexasomes and nucleosomes (Fig. S4C). Moreover, we determined that the unwrapping trajectories of these bigger (H3-H4)_2_ oligomers resemble those of purified hexasomes and exhibit a change in extension of 24.5 ± 1.5 nm centered at 16.9 ± 3.8 pN (n= 14) (Fig. S4D).

### Nucleosome, hexasome, and tetrasome rewrapping trajectories

Unwrapping trajectories of nucleosomes were observed in successive pulling curves during disassembly experiments, indicating that during relaxation full rewrapping had taken place (Fig. 3A). The relaxation curves following the first pulling of nucleosomes are identified as nucleosome rewrapping trajectories only if the subsequent pulling curve displays the low- and high-force transitions. The relaxation trajectories can show one (Fig. 3B), two (Fig. 3C), or three (Fig. 3D) shortening transitions over a broad range of forces (2-15 pN). Having characterized the unwrapping trajectories of tetrasomes and hexasomes, we were able to interpret and assign the relaxation transitions to the rewrapping of specific nucleosomal intermediates.

**Fig 3.**
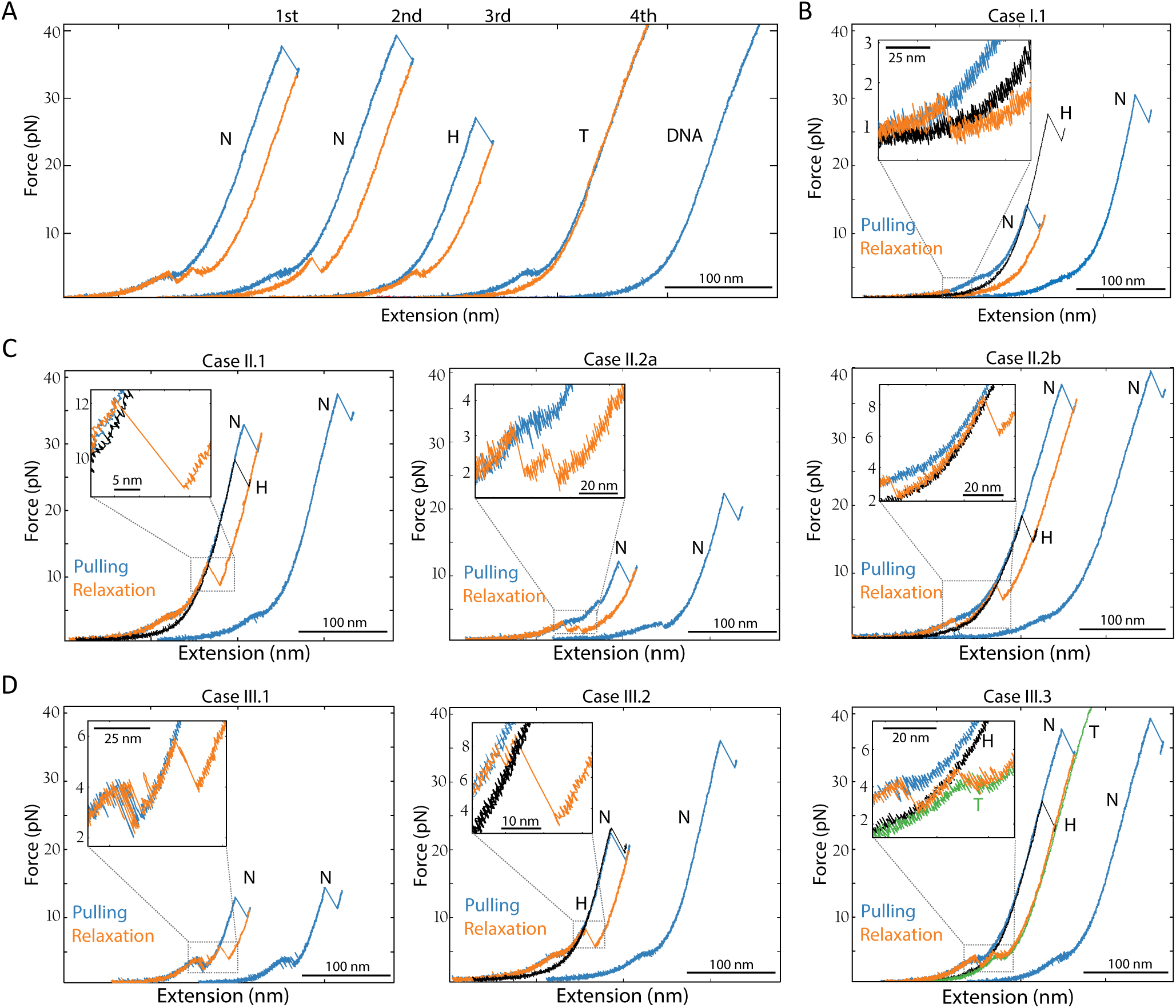
Reversible assembly of nucleosomes. (A) Nucleosome (N) disassembly at 50 mM KOAc by four cycles (1st to 4^th^) of pulling (blue curve) and relaxation (red curve). Pulling and relaxation cycles has been displaced laterally for clarity. The second pulling corresponds to the unwrapping trajectory of a nucleosome, which indicates nucleosome rewrapping along the first relaxation. Hexasome (H) and tetrasome (T) were observed in the pulling curves of the third and fourth cycles, respectively. (B) Nucleosome rewrapping via a single-zip at ~1 pN. Hexasome (H) pulling curve (black curve) was superposed to identify the formation of intermediates. (C) Three types (cases II.1, II.2a, and II2.b) of nucleosome rewrapping via two shortening transitions. See text for description. (D)Three types (cases III.1, III.2, and III3) of nucleosome rewrapping via three shortening transitions. In case II.3, the tetrasome (T; green curve) and hexasome (H; black curve) unwrapping trajectory and was superposed to identify the rewrapping of intermediates. See text for description.

The single shortening transition occurs at a force below the low-force unwrapping transition (Fig. 3B), it is cooperative (zip), and shows a decrease in extension of ~40 nm, corresponding to the rewrapping at once of the DNA unwrapped during the previous low- and high-force transitions (case I). When the relaxation curve displays two shortening transitions, there are three cases. In the first case (case II.1), one of the transitions is a zip that occurs at a force higher than the low-force unwrapping transition (Fig. 3C; left panel); in this case, the rewrapping trajectory reaches the partially unwrap nucleosome pulling curve and it is continued by a second non-cooperative shortening transition that follows the pulling curve all the way to zero force resulting in a fully wrap nucleosome. The zip exhibits a decrease in extension of ~22 nm indicating that it corresponds to the rewrapping of the DNA unwrapped during the previous high-force rip. In the second case (Case II.2), the relaxation curve displays two zips; these can be of variable size, and may occur both below (case II.2a; Fig 3C; middle panel), or one above and one below (case II.2b; Fig 3C; right panel) the low-force unwrapping transition. In these cases, the first zip corresponds to the rewrapping of the hexasome, whereas the second zip corresponds to the full rewrapping of the nucleosome. When the relaxation curve displays three shortening transitions there are three cases. In the first case (case III.1), the relaxation curves display two zips below or near the low-force unwrapping transition followed by a non-cooperative shortening (Fig. 3D; left panel). In the second case (case III.2), the relaxation curves display two zips above the low-force unwrapping transition followed by the non-cooperative shortening (Fig. 3D; middle panel). In both of these cases, the first zip, as before, reaches the hexasome pulling trajectory, and the second corresponds to the formation of the partially unwrapped nucleosome, followed by the full rewrapping of the nucleosome via the non-cooperative transition. In the third case (case III.3), the first transition is non-cooperative, occurs at a force close to the non-cooperative unwrapping transition and it overlaps with the tetrasome unwrapping trajectory; it is followed by two zips of variable size, the first of which reaches the hexasome pulling trajectory and the second represents the formation of the fully wrapped nucleosome (Figure 3D; right panel). Accordingly, nucleosome rewrapping can occur via the sequential rewrapping around tetrasomes and hexasomes, through hexasomes alone, or in a single step without detectable intermediates.

A similar analysis has been performed for the rewrapping of hexasomes (Fig. S5A) and tetrasomes (Fig. S5B) during nucleosome disassembly. Interestingly, in few cases, tetrasomes that have been generated by nucleosome disassembly in which the pulling and relaxation cycles did not exceed ~10 pN by design, we observed pulling curves going back to those characteristics of the unwrapping of hexasomes, displaying a single rip at high force (Fig. S5C). One possible explanation for this observation is that the H2A-H2B heterodimer remained bound to the DNA during unwrapping and eventually re-engage the tetrasome to reform a hexasome.

### Annotation of mechanically-induced transitions by simultaneous detection of force and FRET signals

Ngo et al. used single-molecule FRET experiments of nucleosomes labeled at different positions of the 601 sequence, to show that under tension unwrapping is asymmetric and its directionality is dictated by sequence dependent DNA flexibility (22)). These authors showed that unwrapping begins at low force with the less flexible DNA arm (monitored by a FRET pair labeled ED1), followed by an internal position opposite to the dyad at higher force (monitored by a FRET pair labeled INT) and ending with the unwrapping of the more flexible arm (monitored by FRET pair ED2).

In order to unambiguously assign the force-extension transitions of nucleosome unwrapping with the specific DNA segments involved, we used an optical tweezers instrument with simultaneous single-molecule fluorescence detection capability or “fleezers” (36–38). To that end, we synthetized and purified ED1, INT, and ED2 nucleosomes (labeled with Cy3/Cy5 pairs) (Fig. 4A), which were ligated to 2.5 kb DNA handles to allow their attachment to functionalized polystyrene microbeads (Fig. 4B). We first confirmed the presence of both dyes and the FRET state corresponding to a fully wrapped nucleosome using the confocal scan of the instrument (Fig. 4C). Next, we monitored the co-temporal evolution of force, extension, and fluorescence of individual fluorophores and FRET of ED2 (Fig. 4D), INT (Fig. S6A), and ED1 (Fig. S6B) during pulling and relaxation cycles. We used a symmetric pulling protocol whereby both optical traps were moved simultaneously in order to keep the fluorescently labeled nucleosome at the center of the confocal spot The fluorescence channel shows the anticorrelated changes of Cy5 and Cy3 fluorescence with the corresponding high-to-low and low-to-high FRET transitions during the unwrapping and rewrapping trajectories, respectively. The recovery of FRET in the rewrapping rules out photobleaching of the acceptor dye during the experiments.

**Fig. 4.**
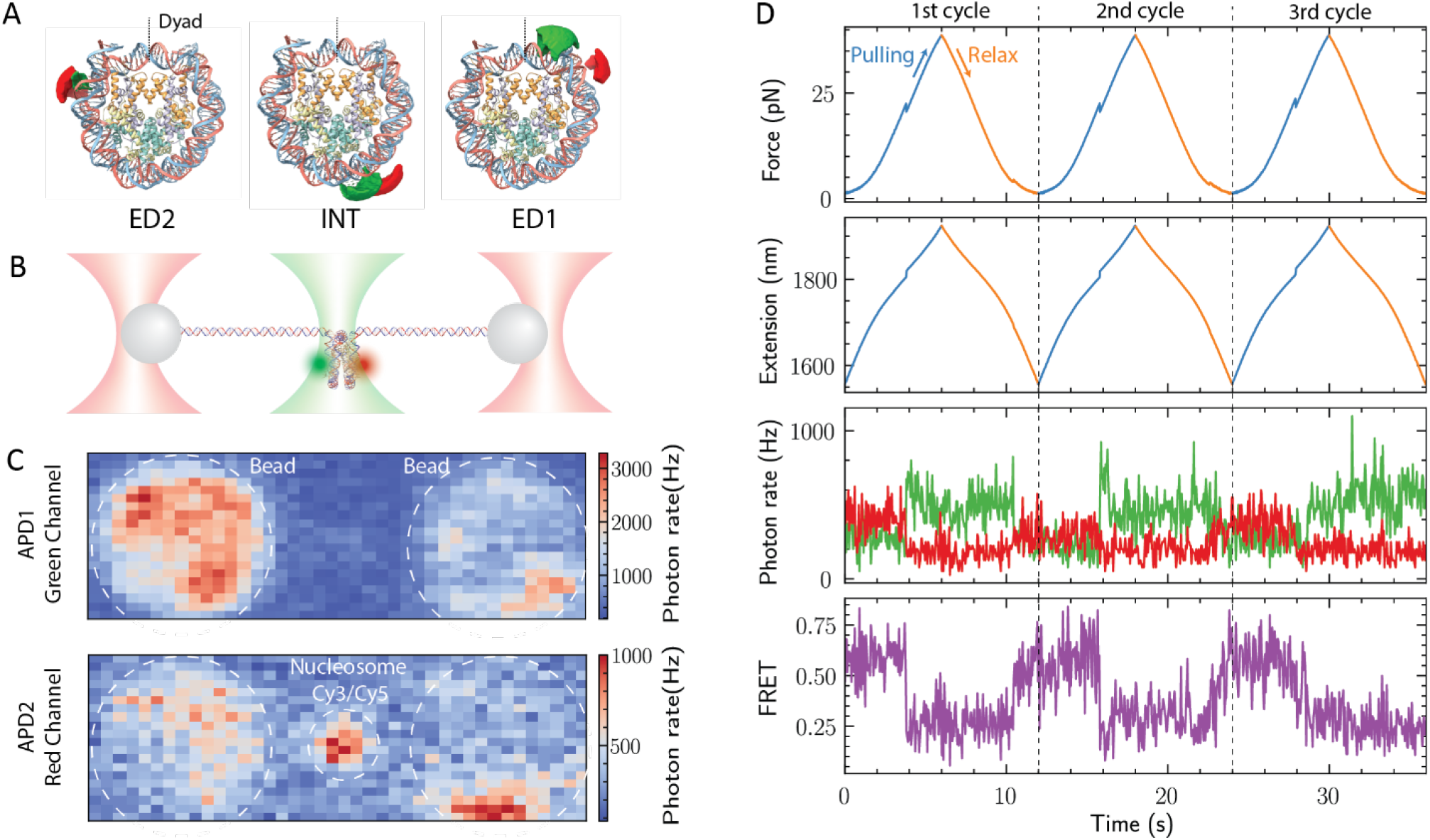
Asymmetric unwrapping of nucleosomes monitored by co-temporal force and fluorescence measurement. (A) Structure of the nucleosome (PDB:6ESF) showing the position of the fluorophores Cy3 (green) and Cy5 (red) in the ED2, INT, and ED1 nucleosomes. (B) Experimental geometry: single Cy3/Cy5 nucleosomes tethered for mechanical manipulation in a “fleezers” setup. (C) Confocal scanning displaying the fluorescence of a single tether of Cy3/Cy5 nucleosome under 532 nm green laser excitation. (D) Simultaneous force, extension, and fluorescence measurements of a single tether of ED2 nucleosome during three pulling and relaxation cycles (separated by the black dashed lines). In force and extension plots, blue and orange traces indicate pulling and relaxation cycles, respectively. Fluorescence channel detects the anticorrelated changes in green and red signals corresponding to changes in FRET. Distinctive and simultaneous transitions in the force, extension, and FRET occur at high-force (~20 pN) due to unwrapping events. The recovery of the FRET signal indicates nucleosome rewrapping.

Detailed comparison of the time course between force-extension and FRET signals of all the molecules tested indicates that, within our temporal resolution (10 ms), for both ED2 (Fig. 5A and B) and INT (Fig. 5C and D) nucleosomes, the FRET change occurs simultaneously with the force-extension transition at high-force, regardless of whether the low-force transition was cooperative or non-cooperative (left and right panels in Fig. 5A and C, respectively). We noted that previous to the transition, the INT nucleosome exhibits larger fluctuations in FRET compared to ED2 nucleosome, a behavior previously described as “gaping” (39). The simultaneous decay of FRET with the high-force rip implies that the changes in end-to-end distance (measured by optical tweezers) and the local DNA changes (as monitored by FRET) report on the same DNA unwrapping process.

**Fig. 5.**
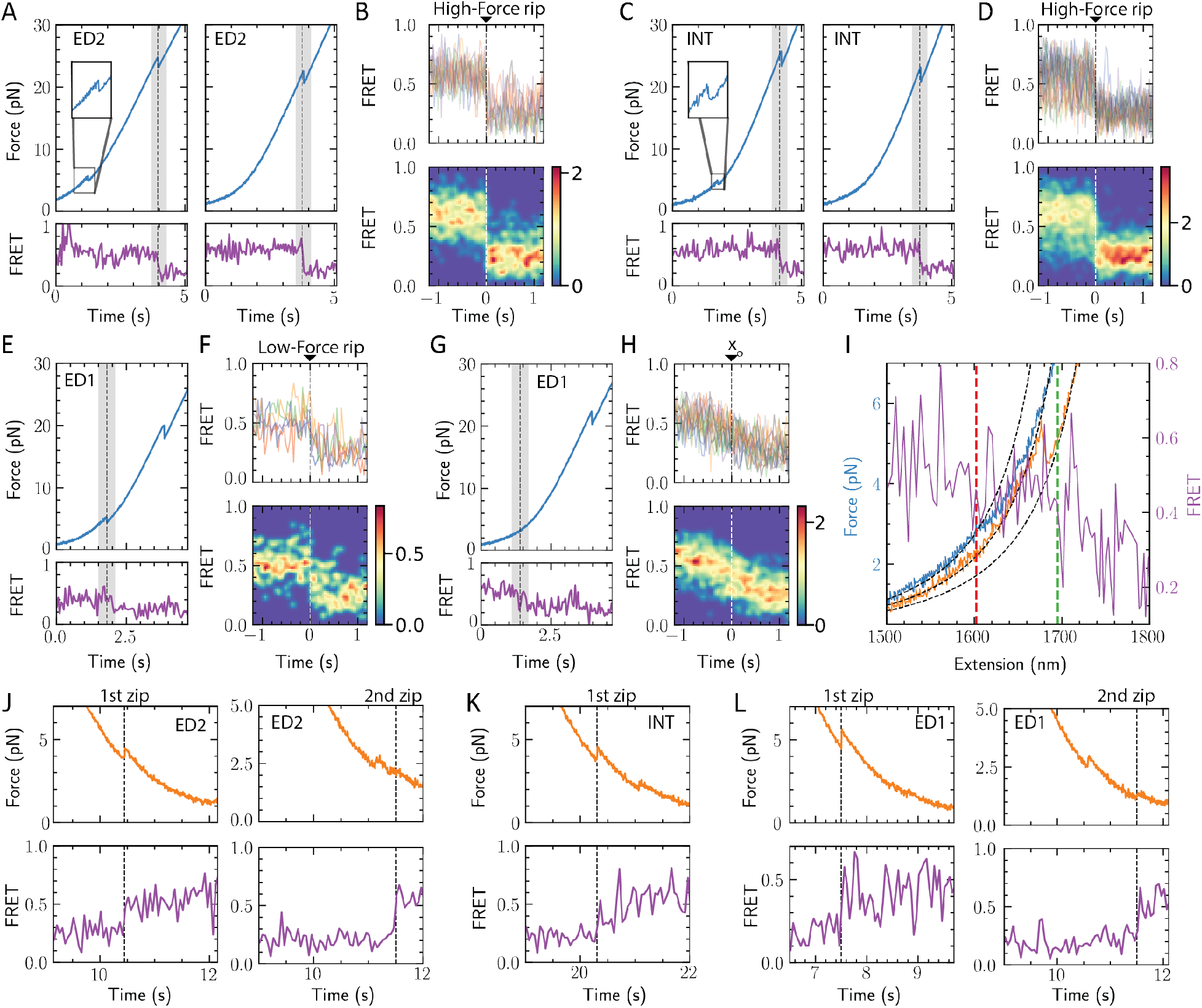
Unwrapping and rewrapping trajectories of FRET nucleosomes. Force pulling trajectory with the corresponding FRET evolution for (A) ED2, and (C) INT nucleosomes exhibiting a cooperative (left panel) or non-cooperative (right panel) force-extension transition at low-force. The shaded grey area highlights the high-force rip. Alignment of FRET transitions for (B) ED2 (n= 12), and (D) INT (n= 23), using the high-force rip as a fiduciary mark (t= 0), shows the simultaneous drop in both FRET and force (top panels displays aligned traces; bottom panels display a 2-D histogram of aligned traces). (E) Force pulling trajectory and FRET evolution of ED1 nucleosome exhibiting a cooperative force-extension transition at low-force (grey area). (F) Alignment of ED1 FRET transitions (n= 5) using the cooperative low-force rip as zero time shows a decrease in FRET as one-step transition. (G) Force pulling trajectory and FRET evolution of ED1 nucleosome exhibiting a non-cooperative force-extension transition at low-force (grey area). (H) Alignment of ED1 FRET transitions (n= 17) exhibiting a non-cooperative force-extension transition at low force. FRET traces were first fitted to the inverted logistic function 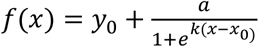, where *x*_0_ was used as zero time for alignment of the traces. (I) Temporal relation between the non-cooperative decrease in FRET as a function of distance (purple trace) and the non-cooperative change in extension of ED1 nucleosomes. The fitting of the worm-like-chain model (dashed black lines) to the low-force section of the nucleosome pulling curve (blue solid line) exhibit a divergence as a consequence of the beginning of the non-cooperative low-force transition (red dashed line). Hexasome relaxation trajectory (orange curve) was included as a reference to validate the worm-like-chain fitting. The shift observed between the non-cooperative force extension transition and the start of FRET decrease (dashed green line) indicates that DNA unwrapping is not the only process occurring at the low-force transition. (J-L) FRET recovery during relaxation (orange curve) indicates rewrapping. (J) ED2 nucleosome rewrapping occurs in the first (n= 3; left panel) or second zip (n= 3; right panel), (K) INT nucleosome rewrapping occurs always in the first zip (n= 21), and (L) ED1 nucleosome rewrapping occurs in the first (n= 3; left panel) or second zip (n= 2; right panel), indicating that nucleosome rewrapping does not exhibit asymmetry.

In contrast, for ED1 nucleosomes we observe two distinct FRET changes at low-force: i) a sharp decrease in FRET, which coincides with the force-extension transition when this occurs cooperatively (Fig. 5E and F), and ii) a gradual FRET decrease that occurs along with the non-cooperative force-extension transition (Fig. 5G and H). We stress that these alignment analyses were performed with a clear force-extension change observed for all cooperative transitions; however, such analysis is compromised for the non-cooperative transitions due to their gradual change in force-extension at low-force. Moreover, we note that unlike unwrappping experiments with shorter DNA handles (Fig. 1), the longer handles required for “fleezers” experiments increase the noise making it difficult to determine the beginning and the end of the non-cooperative force-extension transition. As a result, we lack a fiduciary mark for the alignment of the corresponding FRET traces. Therefore, we fitted the FRET traces to an inverse logistic function and we used the midpoint of this fitting as reference point to align the traces (Fig. 5H). The resulting alignment shows that the decrease in FRET is also a non-cooperative process.

To determine the temporal relation between the non-cooperative decrease in FRET and the non-cooperative change in extension in the optical tweezers channel, we need to locate first the non-cooperative force-extension transition. To this end, we compared the first pulling curve of a nucleosome with one of its subsequent relaxing curves that follows the unwrapping trajectory of a hexasome (Fig. 5I and Fig. S7). The point at which the experimental pulling curve of the nucleosome diverges from its theoretical worm-like chain (red dashed line) determines the start of the non-cooperative transition (see red dotted lines in Fig. 5I and Fig. S7). To assign the relative location of gradual FRET change with respect to the non-cooperative transition, we plotted the change in FRET as a function of extension, taking advantage that the optical tweezers and the fluorescence channels are monitored co-temporally. Comparison of both channels shows that the FRET changes and the non-cooperative force-extension transition do not occur simultaneously (Fig. 5I) with their separation in time varying among different molecules (Fig. S7). This observation indicates the existence of a process that contributes to the change in extension in the optical tweezers channel, without involving DNA unwrapping.

To study nucleosome rewrapping, we analyzed the relaxation trajectories where high-FRET was recovered (Fig. 5J-L). The force-time course shows that nucleosomes rewrap most of the time in two steps and rarely in a single-one at forces below ~6 pN. In the second case it is likely that the second transition was missed due to the low-force at which occurred. While ED2 and ED1 nucleosomes recover their FRET signals during the first or the second zip irrespectively (Fig. 5J and L), INT nucleosomes always recover their high FRET value coincidentally with the first zip (Fig. 5K). Thus, although the unwrapping of nucleosomes occurs asymmetrically and sequentially starting always with the opening of the ED1 arm, the rewrapping can start with either arm in a sequential fashion.

### Real-time observation of H2A-H2B dimer ejection occurs following the high-force transition

As shown above (Fig. 2), the mechanical disruption of nucleosomes leads to the formation of hexasomes and tetrasomes characterized by pulling trajectories displaying a single transition each. We interpret the loss of the low-force transition as reflecting the dissociation of an H2A-H2B dimer. It is, therefore, of interest to establish when during the unwrapping of the nucleosome this dissociation occurs. To this end, we need to correlate the loss of force-extension transition with heterodimer release. Accordingly, we attached a single Cy3 fluorophore via synthetic peptides to the H2A C-terminal tail (H2A-Cy3) using sortase (40, 41). This method also made it possible to purify H2A-Cy3 from the unlabeled H2A. Nucleosomes and hexasomes assembled with H2A-Cy3 were purified and their correct integrity was confirmed by Total Internal Reflection Fluorescence (smTIRF) microscope (nucleosomes, Fig. S8A-C; hexasomes, Fig. S8D-F).

Next, we ligated the H2A-Cy3 nucleosomes and hexasomes to 2.5 kb DNA handles to characterize them using the “fleezers” instrument (Fig. 6A). At low tension we confirmed the presence of two H2A-H2B heterodimers via the two-step photobleaching (Fig. S8G), and the subsequent force-extension pulling trajectory exhibits the low- and high-force transitions of an intact nucleosome (Fig. S8H). Similarly, tethered single hexasomes display only one-step bleaching (Fig. S8I) and their force-extension pulling curves exhibit only the single high-force rip (Fig. S8J). The “fleezers” setup, yielded a lower proportion of two-step bleaching events compared to the smTIRF assay. Therefore, although two-step bleaching events for H2A-Cy3 nucleosomes were observed, single or no fluorescent events were also present in the “fleezers” instrument. We attributed this difference to trapping-laser-induced photobleaching (42). However, the integrity of the nucleosome can be deduced from the force-extension unwrapping trajectory, and the fluorescence signal change can inform us about the precise moment where the H2A-H2B heterodimer is lost during a pulling experiment.

**Fig. 6.**
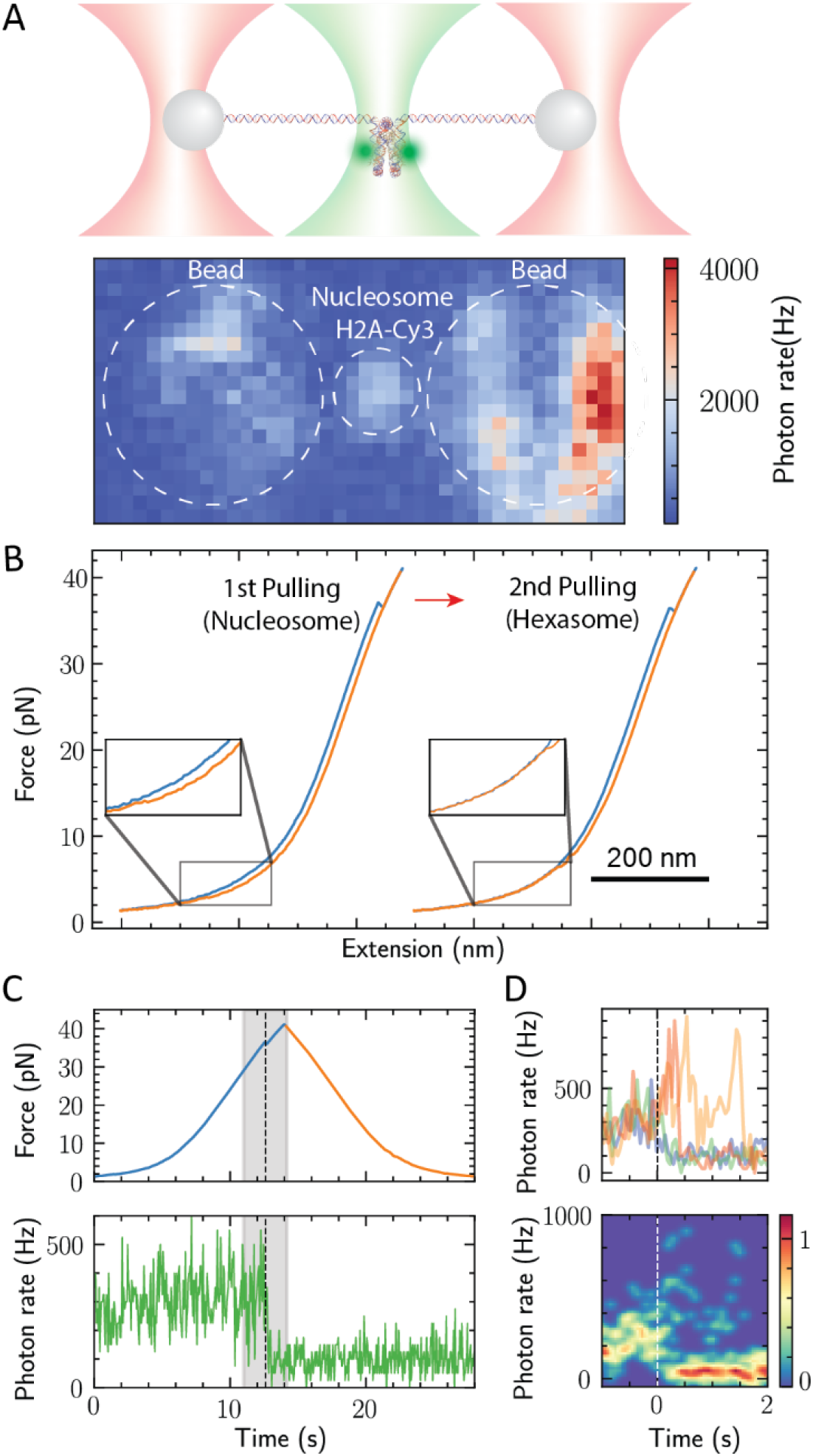
Real-time detection of H2A-H2B heterodimer dissociation during nucleosome mechanical unwrapping. (A) Experimental geometry depicting a single H2A-Cy3 nucleosome tethered for mechanical manipulation in a “fleezers” setup (top panel). Under 532 nm laser excitation, the confocal scan displays the fluorescence of a single H2A-Cy3 nucleosome (bottom panel). (B) H2A-Cy3 nucleosome disassembly by two cycles of pulling (blue curve) and relaxation (orange curve). Pulling/relaxation cycles were artificially separated for illustrative purposes. The lack of the low-force transition in the second pulling curve (inset) indicates the formation of an hexasome after the first unwrapping. (C) Simultaneous time course of the fluorescence and force channels during the first unwrapping/rewrapping cycle monitored in (B). A decrease in the Cy3 fluorescent signal coincides with the high-force rip indicating the loss of a H2A-H2B heterodimer in the second high-force transition. Shaded gray area indicates the regions of the high-force transition. (D) Aligned fluorescent transitions from different nucleosomes (n= 4) using the high-force rip as a fiduciary mark. After the high-force rip, the fluorescent signal fluctuates and increases before dropping at variable times (top panel display aligned traces; bottom panel display a 2-D histogram of aligned traces).

The mechanical unwrapping of Cy3 labeled nucleosomes leads to the formation of hexasomes as shown by the absence of the low-force transition in the second unwrapping trajectory (Fig. 6B). Where does the H2A-H2B heterodimer dissociate from the histone core in the pulling/relaxation trajectories? We observed that the fluorescent signal did not change its average intensity during the unwrapping at the low-force transition (Fig. 6C and S9A-C), confirming that at this force regime the nucleosome did not undergo irreversible conformational changes (17, 21). In contrast, the fluorescence suddenly drops in the region of the high-force rip (Fig. 6C and Fig. S9A-C). These results establish a causal relationship between the extended unwrapping of the nucleosomes occurring at the high-force transition and the loss of integrity of the histone core. Similarly, the disassembly of a hexasome with the loss of the H2A-H2B heterodimer to yield a tetrasome, happens in the region of the high-force rip (Fig. S9D and E). To determine the temporal relationship between Cy3 fluorescence loss and the high-force rip, we aligned the fluorescence of different nucleosomes relative to the time at which the force-extension transition at high-force was detected in each case (Fig. 6D). We observe that, after the high-force rip, the Cy3 fluorescence exhibits large intensity fluctuations before it drops at different time points after the rip (average delay = 113 ± 126 ms, mean and SEM).

### Assignment of changes in extension during mechanical unwrapping of nucleosomes

As shown above, we found an inconsistency between the unwrapping extension observed in the low-force transition with the optical tweezers and that reported by the co-temporal changes in FRET. Indeed, if we were to interpret the extension change of the low force transition as purely reflecting the asymmetrical unwrapping of DNA, it would correspond to ~60 bp of DNA (20.4 nm). This extent of unwrapping should be accompanied by a loss of the FRET signal at the INT position, which is located 28 bp away from the entry point of the weak arm. However, as shown above, this FRET change only occurs during the high-force transition (Fig. 5C). Furthermore, the acceptor dye of the FRET pair closest to INT used by Ngo et al. (ED1.7) ceases to fluoresce at low-force and it is localized 24 bp from the entry point of the weak arm (22). This observation indicates that the low-force transition should correspond to the unwrapping of only ~24-27 bp of DNA or about one half of the outer wrap of the DNA in the crystal structure. This analysis indicates that DNA unwrapping from the histone core is not the only process contributing to the extension observed in the low-force transition with the optical tweezers. Moreover, if only ~27 bp are unwrapped in the low-force transition, there should be ~120 bp left partially wrapped on the nucleosome. However, the change in extension in the high-force transition (~24.5 nm) is significantly shorter than that expected if it involves the complete unwrapping of the nucleosome as previously proposed (17, 21, 31).

The presence of an additional process not involving unwrapping during the non-cooperative low-force transition is confirmed by the change in extension detected in the optical tweezers signal before DNA unwrapping takes place as reported by FRET (Fig. 5I and S7). Such additional process has been hypothesized in the spool model of nucleosome unwrapping, which proposes that as a consequence of its spool geometry a nucleosome must reorient, rotate, and align when subjected to tension (21, 43, 44). However, although the spool model addresses the energetics barriers for DNA unwrapping, it did not consider the contribution of nucleosome reorientation to the observed change in extension.

We propose that at zero tension, nucleosome DNA linkers cross and emerge in opposite directions (Fig. 7A; inset 1). The increase in tension separates and bends the DNA linkers, and causes the gradual rotation of the nucleosome, marking the beginning of the non-cooperative low-force transition (Fig. 7A; inset 2). Once rotation has begun, ~27 bp of DNA unwrap progressively until the nucleosome gets aligned with the applied force. To estimate the contribution of nucleosome reorientation on the observed change in extension, we developed a mechanical unwrapping model that simulates the rotational motion of a nucleosome by aligning the torque exerted at the entry and exit DNA along the pulling force direction and the unwrapping transitions (Supplementary Movie).

**Fig. 7.**
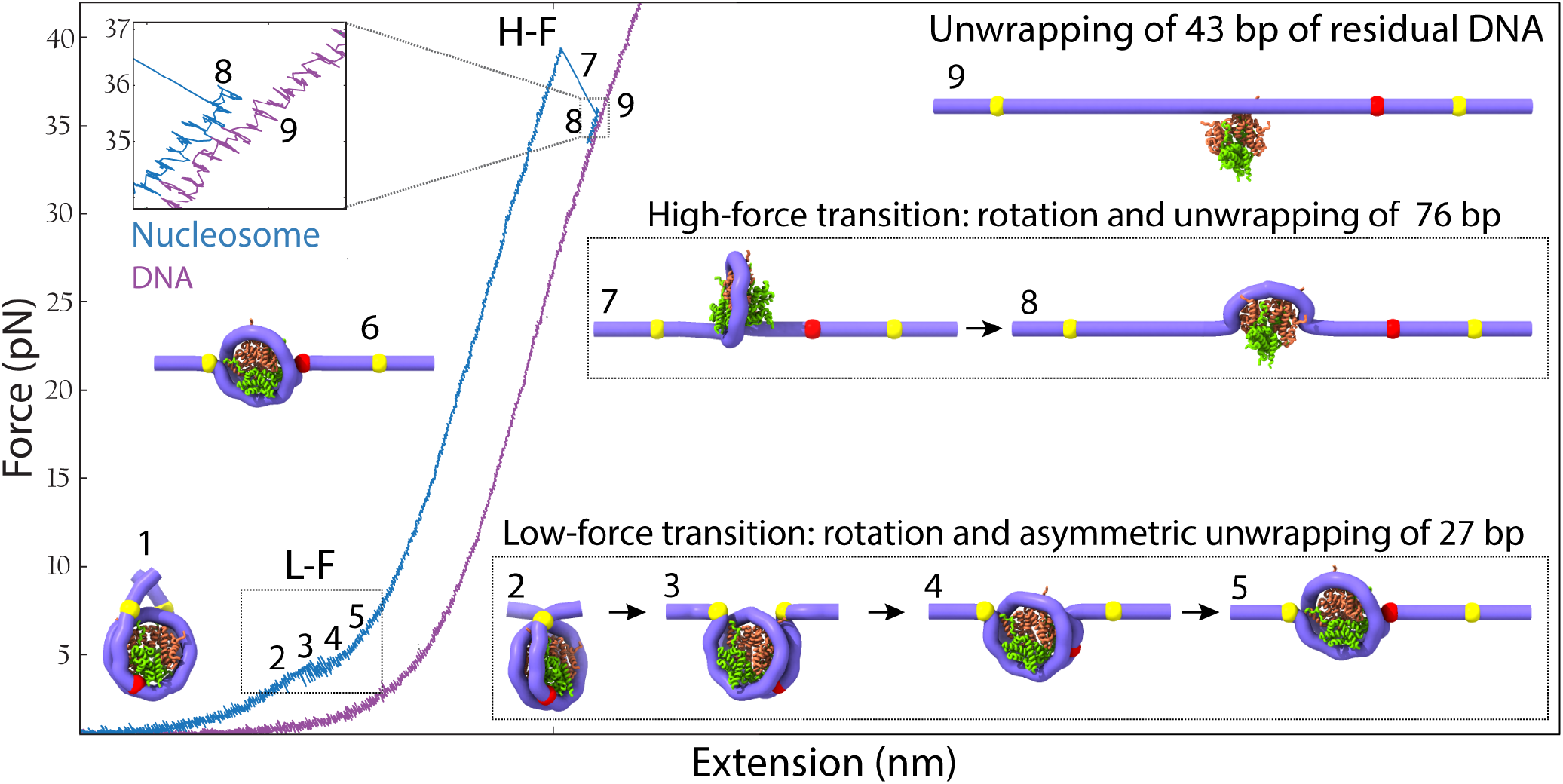
Annotation of the structural changes corresponding to the force-extension transitions during the mechanical unwrapping of a nucleosome. Upon the application of the external force, the nucleosome arms at zero force (inset 1) align along the pulling axis (inset 2). During the non-cooperative low force-transition (~4 pN, L-F, insets 2-5), the nucleosome first rotates without unwrapping keeping the DNA entry and exit points in contact with the core particle (yellow marks). At this point, it begins to unwrap asymmetrically and progressively by 27 bp (red mark) corresponding to the detachment of the distal or weak DNA arm (insets 4-5). This last conformation (inset 5) is maintained along the pulling curve (inset 6) until it reaches the high-force transition (H-F). During the high-force transition, nucleosome simultaneously rotates by 180° and unwraps ~76 bp in a single-step observed as a rip (inset 7-8). At the end of the high-force transition, ~43 bp remained wrapped at the histone core (inset 8). Around 13 bp are further unwrapped non-cooperatively at the end of the high force transition, leaving about 30 bp wrapped around the nucleosome which are experimentally observed as a difference in extension (top left inset) of the unwrapped nucleosome compared to bare DNA (purple curve).

Initial rotation of the nucleosome before unwrapping contributes an increase in extension of 8.1 nm (Fig. 7A; inset 3). At this point an increase in end-to-end distance of ~8.5 nm occurs associated with the nucleosome unwrapping asymmetrically by ~27 bp (calculated from the WLC model at a force of 4 pN) plus a residual rotation of ~3.2 nm taking place during the unwrapping (insets 4 and 5). This analysis yields a total change of ~20 nm at the end of the low-force transition. When the low-force transition occurs cooperatively, the nucleosome rotation and unwrapping occur simultaneously yielding an experimental change in extension of 20.4 nm ± 1.5 nm (Table S1). When the transition occurs non-cooperatively, there is a larger uncertainty in the change in extension and we obtained a value of ~19 nm. At the end of the low-force transition, ~120 bp remain wrapped in the nucleosome.

In the high-force transition (Fig. 7A), a partially unwrapped and force-aligned nucleosome (inset 6) continues to rotate by ~180° (and therefore the rotation does no contribute to a change in extension; inset 7 and 8) as it further unwraps (45) to yield an experimental change in extension of 24.5 ± 1.6 nm in a single step (Table S2). The sharp drop in FRET at the entry (ED2) and exit (INT) DNA points coincides with the rip (Fig. 5A-D), indicating that, unlike what is observed at low-force, during the high-force transition DNA unwraps from both ends simultaneously. Using our mechanical unwrapping model, the ~24.5 nm change in extension observed in the high-force transition corresponds to the unwrapping of ~76 bp of DNA. Accordingly, ~43 bp remained wrapped at the end of the high-force transition.

This analysis predicts that a difference in end-to-end distance of 6.2 nm should be observed between the nucleosome at the end of the high-force transition and a bare DNA molecule of the same total length (inset 9; see also Supplementary movie). In support of the mechanical unwrapping model, we developed a geometrical model of nucleosome unwrapping which treats the nucleosome as a spherical particle (Fig. S10 and Supporting information for details). Applying this model to the high-force transition, we determined that the 24.5 nm of change in extension observed during the high-force transition should correspond to ~74 ± 5 bp of DNA, in close agreement with the mechanical unwrapping model. However, experimentally we observe a difference in end-to-end distance between the nucleosome pulling curve after the high-force transition and the corresponding pulling curve of bare DNA of ~2.3 nm (Fig. 2E). Using the geometrical model, we estimate that this difference in extension should correspond to 30 ± 3 bp remaining wrapped (and not 43 bp) after the high-force transition. Thus, we propose that during or after the high-force transition, 13 ± 3 bp unwrap non-cooperatively, a process not directly resolvable with our methodology. Interestingly, 30 bp distributed symmetrically around the dyad has been experimentally determined to be the strongest region of histone-DNA interaction (46).

A similar analysis indicates that the hexasome does not display a low-force transition because has lost the wrapping of one of the arms and the resulting structure is aligned so that the application of force does not generate a change in extension due to rotation. In the case of the tetrasome, the remaining difference of ~6 nm with respect to bare DNA (Fig. 2E) indicates that ~40 bp remain wrapped at the end of the low-force transition, which must involve the unwrapping of ~30 bp and a contribution due to rotation.

## Discussion

Although mechanical manipulation of nucleosomes has been studied for around 20 years, the elucidation of the molecular changes occurring during unwrapping and rewrapping remained controversial. Using FRET in a co-temporal fashion with optical tweezers, we have been able to assign for the first time the observed force-extension transitions to the unwrapping and rewrapping of specific regions in the nucleosome. We find that during unwrapping, nucleosomes can disassemble stochastically into hexasomes and tetrasomes whose force-extension trajectories we establish. We propose a model of nucleosome unwrapping that explain previous discrepancies related to the directionality and extension of DNA unwrapping (17, 21, 22). Furthermore, we show that the assignment of the changes in extension accompanying the unwrapping transitions must include the contribution of the force-induced rotation of the nucleosome as it unwraps.

As previously determined, the entry and exit arms of the 601 NPS sequence possess different mechanical flexibility, which results in the asymmetric unwrapping of the two arms (22). The wrapping of the DNA around the histone core is energetically costly due to the large persistence length of DNA (~50 nm). This cost must be paid by the binding energy between the DNA and the histones. The mechanically stiffer segment corresponds to the DNA weak arm because more of its binding energy to the histones must be used relative to the strong arm to wrap it around the core. As a result, the weak arm unwraps first under mechanical manipulation. Accordingly, the asymmetric unwrapping results from both the differential flexibility of the DNA arms and their interactions with the histone core. When DNA sequence flexibility is symmetrized, either arm can then open before the other because under tension the unwrapping involves the thermally-induced crossing of a barrier, a stochastic process. It has been shown that once one of the arms opens, a conformational change and structural re-accommodation of the nucleosome takes place. Indeed, once ED1 arm unwraps, it triggers conformational changes in different regions of the octamer that bring to closer proximity the opposite ED2 arm and mechanically stabilizes it (47).

The unwrapping of a nucleosome arm at low forces indicates that the interactions involved in maintaining its wrapped structure are relative weak. Indeed, single-molecule unzipping experiments found that this is the case (Hall et al. 2009). Recently, molecular dynamic simulations identified a region in H3 (H3-latch) that in combination with its N-terminal tail interacts with both the inner and outer DNA wrap, stabilizing the DNA arm at the superhelical location (SHL) +7 and keeping it wrapped (Armeev et al. 2021). The limited extent of the low-force transition (~27 bp) probably reflects the strength of the subsequent interactions involving those between the α1 helices and tails of H2A and H2B with the SHL ±4 and with the SHL ±5 (where the INT FRET pair is located; Fig. S11A and B). These regions exhibit a local concentration of positive charges that interact with the negative backbone of the DNA as revealed by calculations of the nucleosome electrostatic surface potential using the Poisson-Boltzmann equation (Fig. S11C and D) (48).

Our observation of a high-force transition confirms the fact that when one of the outer arms of the nucleosome is unwrapped at forces below 5 pN (low-force transition), the remaining nucleosome structure reaccommodates, and the interaction of the opposite arm with the histone core is energetically stabilized so that it will only unwrap at a mean force above 25 pN (high-force transition). This interpretation also explains the unwrapping trajectory of the hexasome: removal of one heterodimer is similar (but not identical) to the mechanical opening of one arm of the nucleosome. The loss of the dimer leads to the stabilization of the DNA arm wrapped around the remaining heterodimer. As a result, the hexasome trajectory lacks the low-force and displays only a high-force transition somewhat weakened relative to that of the nucleosome trajectory (occurring at a mean force of ~25 pN instead of ~31 pN, respectively. Fig. 2B and E). Part of the reaccommodation and stabilization of the nucleosome harboring an unwrapped arm or of the hexasome missing a heterodimer could originate in a reduced electrostatic repulsion between wrapped DNA segments in the nucleosome that accompanies the first unwrapping event ((43). When at this high force, the most external points of contact between the DNA and the histone finally ruptures, most interactions following those points are unable to contain the propagation of the unwrapping front, giving rise to the cooperative nature of the high-force transition. At this point only interactions between SHL +/-1 and SHL +/-2 with α1 helices and tails of H4 and H3 remain (Fig. 7C, left panel).

We find that while the unwrapping process is asymmetric and starts at the weak nucleosomal DNA arm, the rewrapping starts always at the INT segment followed by either the weak or the strong arm indistinctively (Fig. 5J and L). However, the rewrapped nucleosome recovers its unwrapping asymmetry in the next pulling cycle (Fig. 4D and Fig. S6).

It has been shown that Pol II is not strong enough to separate the DNA wrapped around the nucleosome. Instead, it advances through the nucleosome by rectifying the fluctuations of the latter (49). Structural and biochemical studies have shown that Pol II transcription exhibits major pauses at nucleosomal positions SHL ±5 and ±1 independently of the nucleosomal arm from which transcription started (50–52). Taken together, these observations can be rationalized in the context of our nucleosome unwrapping model involving the low-and high-force transitions. First, Pol II exhibits a weak pause at SHL −7 and awaits spontaneous fluctuations that unwrap ~27 bp of DNA (low-force transition) allowing Pol II entry into the nucleosome until it reaches SHL −5. Second, Pol II paused at the SHL −5 awaits larger thermal fluctuations to overcome strong histone-DNA interactions associated with the high-force transition, thus constituting a major pausing site. We note that it is possible that the forces needed to break-up these interactions by thermal fluctuations are lower compared to the forces estimated in our experiments due to our pulling geometry (44, 45) and the bulkiness of Pol II. During transcription, which allows invasion of SHL −1 without breaking the interactions of the distal heterodimer with SHL +5 (51, 52), these interactions are not broken at the same time.

DNA unwrapping at the low-force transition exposes partially one H2A-H2B heterodimer but it does not lead to its dissociation, as confirmed by our “fleezers” assay (Fig. 6) and further supported by: i) the reversible nature of the low-force transition (17, 21), ii) the structure of the partially unwrapped (~25–30 bp) nucleosomes observed by cryo-EM (47), and iii) the exposed H2A-H2B observed by cryo-EM in nucleosomal transcription complexes stalled at SHL −5 (51, 52). Our experiments show, however, that H2A-H2B heterodimer dissociation can occur after the high-force transition in which several DNA-histone interactions are broken. These contacts are partially electrostatic, as indicated by the decrease in the rupture force observed with increasing ionic strength (Table S2). This result is in agreement with single-molecule FRET measurements of salt-induced nucleosome disassembly showing that H2A-H2B dimer dissociation requires extensive DNA unwrapping (53). Furthermore, the rupture of interactions between H2A-H2B and SHL +5 is required for the loss of the heterodimer, but not sufficient. Indeed, we note that after the first pulling and relaxation cycle, hexasomes or tetrasomes are generated only in ~28% of the time in each case, with the remaining (40%) corresponding to the nucleosome retaining its integrity (Table S3). These observations suggest that histone-histone interactions can preserve the H2A-H2B dimer attached to the histone core after DNA unwrapping. Although it remains unclear in our assay which one of the two H2A-H2B heterodimers (proximal or distal) is dissociated during the high-force transition, we speculate that the distal heterodimer (associated with the more rigid ED1 DNA arm) is the first to dissociate based on the preferential assembly of hexasomes containing this proximal heterodimer (54, 55).

In summary, we identify the DNA regions involved with each transition observed in the mechanical unwrapping/rewrapping trajectories of nucleosomes, as well as of the hexasomes, and tetrasomes generated during the stochastic disassembly of the former. The characterization of their mechanical properties will make it possible to identify the subnucleosomal particles generated by the action of polymerases, remodelers, and histone chaperones, in future force spectroscopy studies.

## Materials and Methods

A detailed description of the experimental procedures including protein and DNA preparation, instrument setup, data acquisition and analysis, and the mechanical and the geometrical model for nucleosome unwrapping is included in SI Materials and Methods.

## Acknowledgments

We thank Professor Geeta Narlikar for providing the expression vectors of *X. laevis* histones. We thank Dr. Simon Pöpsel and Professor Eva Nogales for their help with preparative electrophoresis purification. This research was supported by the Nanomachine program (KC1203) funded by the Office of Basic Energy Sciences of the U.S. Department of Energy (DOE) contract no. DE-AC02-05CH11231 (C.B.), and by the National Institute of Health grant R01GM032543. C.B. is a Howard Hughes Medical Institute Investigator.

## Author contributions

C.D.C., C.C.-C, and C.B. conceived the project and designed the research. C.D.C., C.C.-C., R.P.S., J.P.C. and C.B. wrote the paper. C.D.C., A.CH., and M.V. prepared and purified DNA templates, nucleosomes, hexasomes, and tetrasomes. C.C.-C., C.D.C., and E.CH., performed single-molecule fluorescence experiments and C.C.-C analyze FRET data. B.O. collected and analyzed AFM images. R.P.S., J.H.K., M.Z., C.D.C., and J.P.C. performed nucleosome pulling experiments and. R.P.S. and J.P.C analyze the data. M.Z. developed the nucleosome rotational model.

## DECLARATION OF INTERESTS

The authors have declared that no competing interests exist.

## Notes

### Competing Interest Statement

The authors have declared no competing interest.

