## Supporting Information for "Assignment of structural transitions during mechanical unwrapping of nucleosomes and their disassembly products"

### SI Materials and Methods

#### Protein expression and purification

Recombinant *Xenopus laevis*, human, and yeast histones H2A, H2B, H3, and H4 were expressed in *E. coli* BL21(DE3) and purified from inclusion bodies as previously described (1). Recombinant His-tagged sortase 5M in pET30b vector was purified as previously described with some modifications (2). Sortase vector was transformed into *E. coli* BL21(DE3) and bacterial cells were grown while shaking at 37˚C in DYT media supplemented with kanamycin and chloramphenicol. Sortase expression was induced at an OD600 of ~0.6 by adding Isopropyl b-D-1-thiogalactopyranoside (IPTG) at a final concentration of 1 mM. After 3 h of induction at 30˚C, cell cultures were pelleted, washed, resuspended in wash buffer (50 mM Tris-HCl pH 7.5; 300 mM KCl; 0.1 mM EDTA; 40 mM imidazole; and protease inhibitors (Roche)), sonicated, and centrifuged at 25,000 x g for 30 min. Supernatant was loaded into a HisTrap SP column previously equilibrated with wash buffer, and sortase was eluted with a linear gradient of imidazole (40-1,000 mM). Fractions were checked using 15% SDS-PAGE, and fractions containing the sortase were pooled and dialyzed using 20mM Potassium phosphate, pH 6.0, 1 mM 2-mercaptoethanol (BME). The dialyzed sortase was loaded on a HiTrap SP cationic exchange column (previously equilibrated with 20mM Potassium phosphate, pH 6.0, 1 mM BME), and sortase was eluted with a gradient of 0-250 mM KCl. Fractions containing the sortase were pooled, concentrated by centrifugation using a 10K Amicon Ultra-15, and loaded on a Sephacryl S100 column previously equilibrated with 25 mM Tris-HCl, pH 7.5, 150 mM KCl, and 1 mM DTT. Purification was checked using 15% SDS-PAGE, and fractions containing pure sortase were pooled and concentrated by centrifugation. Sortase concentration was calculated using an extinction coefficient of 14,440 $M^{-1}cm^{-1}$, and the concentrated sortase was combined with glycerol (30% final) then stored at -80˚C.

#### DNA templates

The nucleosomal DNA used in this work corresponded to the 601 and the 5S nucleosome positioning sequences (NPS). The 601 mononucleosome DNA consists of one repeat of the 601 NPS flanked to the left by 100 bp DNA and to the right by 50 bp DNA (100W50). To produce large amounts of the DNA template, 100W50 was amplified by PCR from the PGEM 601 vector, and twelve copies of 100W50 were cloned back into the PGEM 601 vector using primers containing the restriction recognition sites for enzymes *KpnI*, *BamHI*, and *BglII* (NEB), as previously described (3). Primers also included the recognition site for *BsaI* (NEB) to excise the 601-NPS repeats from the vector and to ligate to DNA handles in nucleosome pulling experiments. The 5S nucleosomal DNA consists of one repeat of the 5S-NPS flanked to the left and right by 20 bp DNA. 5S DNA was designed with shorter DNA linkers to prevent mispositioning of the histone core. 5S template was amplified by PCR from a gBlock that codifies the 5S sequence of sea urchin (GenBank: V00645.1), and one copy was cloned into the PGEM 601 using the *BsaI* enzyme. Plasmids containing the 12 repeats of 100W50 or the single copy of 5S were grown in *dam⁻/dcm⁻ E. coli* (NEB), purified by maxiprep, and excised by restriction with *BsaI*. 100W50 and 5S templates were purified from the vector backbone by 5% preparative acrylamide electrophoresis using a Model 491 Prep Cell (Bio-Rad). For smTIRF imaging, 100W50 template was synthetized by PCR from the PGEM 601 vector using a forward primer with a 5' biotin modification (IDT) for surface attachment and was purified by 5% preparative acrylamide electrophoresis.

#### Synthesis of fluorescently labeled 601 NPS

We used PCR to generate ED1, INT, and ED2 fluorescent DNA templates. The primers used are based on published work but were modified to incorporate flanking *BsaI* restriction sites for ligation with handles and were PAGE-purified. To attach the fluorescent labels to these primers, Cy3-NHS and Cy5-NHS (Lumiprobe) were conjugated to the forward and reverse primers, respectively. Our labeling efficiency was >90%. Labeled primers were used to amplify the 601 NPS from the PGEM 601 vector using the Taq DNA polymerase (NEB). Fluorescently labeled DNA templates were concentrated by centrifugation using a 10K Amicon Ultra-15 filter, and the PCR buffer was exchanged with 10 mM Tris-HCl pH 8.0. DNA templates were digested with *BsaI* and purified by 5% preparative acrylamide electrophoresis. Fractions containing the DNA template were pooled, concentrated by centrifugation, and stored in the dark for later nucleosome reconstitution.

#### H2A fluorescent labeling using sortase

**Preparation of GGGK(Cy3) peptide.** An N-terminal Fmoc protected peptide (Fmoc-GGGK, Genscript) was reacted overnight and at room temperature with a 5-fold excess of Cy3-NHS

(Lumiprobe) in coupling buffer (0.2 mM NaHCO3 pH 8.5, 10% DMSO). The reaction was then treated with 50% ethanolamine to remove with Fmoc group and quench remaining Cy3-NHS. Non-polar by-products were removed by solvent-extraction using hexanes. The aqueous pink solution containing the product GGGK(Cy3) was then purified using a C18 column (Sep-Pak tC18 3 cc Vac Cartridge, 200 mg Sorbent per Cartridge, 37-55 µm, Waters). The purified GGGK(Cy3) was rotavaporated, lyophilized, redissolved in 100% DMSO, and stored at -20˚C.

**Expression and labeling of H2A-Sortag with GGGK(Cy3).** The C-terminal of the *X. laevis* H2A gene was tagged using PCR primers carrying the sortase recognition motif (LPETGG) followed by six histidines. H2A-LPETGG-6x-His was expressed and purified following the regular histone protocol (3). The transpeptidase reaction, containing 10 µM sortase 5M, 250 µM H2A-LPETGG-6x-His, and 500 µM GGGK-Cy3 peptide, was performed in 300 mM Tris-HCl, pH 7.5 and 300 mM NaCl. Reaction was incubated at room temperature for one hour, then diluted 10 times in 50 mM Tris-HCl, pH 8.0, 300 mM NaCl, and 20 mM imidazol. Non-labeled H2A-LPETGG-6x-His was removed from reaction using Dynabeads (Invitrogen) that were equilibrated with 50 mM Tris-HCl pH 8,0, 300 mM NaCl, and 20 mM imidazole. The reaction was subsequently dialyzed against 20 mM Tris-Cl pH 7.5, 7 M GdmCl, and 10 mM DTT to remove free peptides and inactivate the sortase. Labeled H2A-LPETGGG-CY3 was concentrated by centrifugation using a 10K Amicon Ultra-15 for histone octamer reconstitution.

**Octamer reconstitution**

The synthesis of the different histone octamers used in this study was performed as previously described (1). Individual histones were dissolved in unfolding buffer (20 mM Tris-HCl buffer pH 7.5, 7 M guanidinium hydrochloride, 10 mM DTT), and H2A, H2B, H3, and H4 histones were combined at stoichiometric ratios of 1.2:1.2:1:1, respectively, at a final concentration of 1 mg/mL. In the case of the Cy3-H2A labeled octamer, H2A was replaced by H2A-Cy3 which was already in unfolding buffer. Unfolded histones were dialyzed for 12 h, using a 3.5 kDa dialysis membrane, against 500 mL of refolding buffer (10 mM Tris-HClbuffer pH 7.5, 2 M NaCl, 1 mM EDTA, 5 mM BME). This dialysis was repeated four times. Refolded histone octamer was concentrated to ~0.5 mL using a 10 kDa Ultra-15 membrane filter (Milipore) and fractionated using the gel filtration column Superdex 200 Increase 10/300 GL (Cytiva) equilibrated with refolding buffer. Fractions were analyzed by 15% SDS-PAGE and AcquaStain protein staining (Bulldog Bio), then fractions containing the octamer were pooled and concentrated to ~10 mg/mL by centrifugation (30 kDa Ultra-15 (Milipore)). Aliquots were flash-frozen and stored at -80˚C for subsequent use. The synthesis of the $(H3\text{-}H4)_2$ tetramer followed the same procedure utilized for octamer reconstitution, but H3 and H4 were combined at a ratio of 1:1.

**Nucleosome, hexasome, and tetrasome assembly and purification**

Histone octamers and 100W50 or 5S DNA templates were combined at a ratio of 1:1, respectively, in 500 µL of high-salt buffer (10 mM Tris-HCl pH 8.0, 2 M NaCl, 1 mM EDTA, 0.5 mM DTT, and 1 mM PMSF) at a final concentration of 100 ng/uL of DNA. Assembly solutions were dialyzed against 500 mL of high-salt buffer for 1 h at 4˚C, followed by a lineal gradient dialysis against 2 L of low-salt buffer (10 mM Tris-HCl pH 8.0, 1 mM EDTA, 0.5 mM DTT, 1 mM PMSF) using a peristaltic pump with a 0.8 mL/min flow rate and continuous stirring. A final dialysis of 3 h in 500 mL of low salt buffer was done to reduce the residual NaCl concentration, and the nucleosome reconstitution was checked by 5% acrylamide native electrophoresis using 0.2X TBE (Tris-borate-EDTA) as a running buffer. *X. laevis* tetrasomes were assembled by combining $(H3\text{-}H4)_2$ tetramers with 100W50 DNA at increasing ratios (1:1.0/1.4/1.8/2.2/2.6), and following the procedure described for nucleosome assembly. Nucleosomes were separated from hexasomes and bare DNA by 5% preparative acrylamide (59:1 acrylamide:bisacrylamide) electrophoresis using a Model 491 Prep Cell (Bio-Rad). Tetrasome purification was performed using 4% acrylamide. For all purifications, the Prep Cell was run at 6 W, and 0.9 mL fractions were collected in 10 mM Tris-HCl pH 8.0, 1 mM EDTA, 1 mM DTT at a flow rate 0.3 mL/min. Purifications were checked by 5% acrylamide native electrophoresis, and sets of fractions containing nucleosomes, hexasomes, or tetrasomes were separately concentrated by centrifugation using 100K Amicon Ultra filters (Millipore). After concentration, samples were dialyzed against HE buffer (20 mM HEPES-NaOH pH 7.5, 1 mM EDTA, 1 mM DTT) and stored at 4˚C.

**Atomic force microscopy (AFM) imaging**

Nucleosomes, hexasomes, and tetrasomes were diluted to 20 nM and crosslinked with 1% formaldehyde for 1 h at room temperature. Crosslinking reactions were quenched by adding 20 mM glycine for 10 min at room temperature, dialyzed against 20 mM HEPES-NaOH pH 7.5, 1 mM EDTA, 1 mM DTT, and centrifugated at 20,000 x g for 10 min to remove aggregates. Crosslinked samples were diluted in 10 mM MOPS pH 7.0 and 5 mM $MgCl_2$ to concentrations of 3.0 nM, 3.5 nM and 4.2 nM for nucleosomes, hexasomes, and tetrasomes respectively. Three to five microliters of the solution were deposited onto freshly cleaved bare mica V1 (Ted Pella Inc.) and incubated for two to five minutes. Upon adsorption, the samples were gently rinsed with Milli-Q water and dried under a nitrogen stream. AFM micrographs were taken with a MultiMode NanoScope 8 atomic force microscope (Bruker Co.) equipped with a vertical engagement scanner E. The samples were excited at their resonance frequency (280-350 kHz) with free amplitudes (Ao) of 2-10 nm and imaged in tapping mode using silicon cantilevers (Nanosensors). The image amplitude (set point As) and A0 ratio (As/A0) was kept at ~0.8 in a repulsive tip-sample interaction regime, and phase oscillations were no greater than ±5 degrees. The surface was rastered following the fast scan axis (x) at rates of 2 Hz, capturing the retrace line to reconstruct the AFM micrographs. All samples were scanned at room temperature in air at a relative humidity of 30%.

**Single-molecule total internal reflection Fluorescence (smTIRF)**

**Surface passivation and sample assembly.** Fluorescent nucleosomes and hexasomes assembled on the 5' biotinylated 100W50 template were attached to functionalized glass slides for smTIRF imaging. Slides and coverslips were prepared using a mixture of mPEG-SVA and Biotin-PEG-SVA (Laysan Bio) following a previously described protocol (4). The day of the experiment, PEGylated slides and coverslips were incubated with MS(PEG)4 Methyl-PEG-NHS-Ester (Thermo Scientific) for 1 h, washed with Milli-Q water, and dried with nitrogen, and a microfluidic chamber was assembled using the PEG-coverslip and PEG-slide. The channel formed between the surfaces was blocked for 1 h by the addition of 1 mg/mL of Acetylated BSA (AcBSA, Invitrogen) and 1 mg/mL tRNA (Invitrogen) in 50 mM Tris-HCl pH 8.0, 50 mM KCl, and 1 mM $MgCl_2$, followed by incubation for 10 min with 50 mM Tris-HCl pH 8.0, 50 mM KCl, 1 mM $MgCl_2$, 0.2 mg/mL AcBSA, and 0.2 mg/mL NeutrAvidin (Thermo Scientific). Excess NeutrAvidin was washed out with 50 mM Tris-HCl pH 8.0, 50 mM KCl, 1 mM $MgCl_2$, 0.2 mg/mL AcBSA. Nucleosomes or hexasomes were diluted to 12.5 pM in dilution buffer (10 mM Tris-HCl pH 8.0, 50 mM KCL, 1 mM $MgCl_2$, 0.02% NP-40) and added to the visualization channel and incubated for 10 min. The visualization channel was washed with dilution buffer to remove non-bound nucleosomes or hexasomes, then filled with imaging buffer (50 mM Tris-HCl pH 8.0, 50 mM KCL, 5 mM $MgCl_2$, 0.5 mg/mL of AcBSA, 0.5 mg/mL tRNA, 0.02% NP-40, 1% glucose, 2 mM TROLOX (SIGMA), 30 μg/mL Glucose Oxidase (SIGMA), 40 μg/mL Catalase (SIGMA).

**smTIRF microscopy.** A custom-built objective-type, wide-field total internal reflection fluorescence (TIRF) microscope was used for single-molecule fluorescence resonance energy transfer (sm-FRET) measurements. An oil-immersion 100x objective lens (Olympus 100× UPlansApo, N.A. 1.4) was used with a diode-pumped 532-nm laser (75 mW; CrystaLaser) for excitation. Fluorescence emission was detected using a EMCCD camera (IXon EM+ 897; Andor). The illuminated area on each channel was ~50 × 25 μm with ~110 nm pixel widths. Images were recorded using an EMCCD Gain of 100 with an integration time of 100 ms. Images were acquired within 1 hour of using the imaging buffer to account for buffer aging (depletion of glucose and buffer acidification).

**Analysis of the smTIRF data.** Single-molecule TIRF data was analyzed using the software iSMS (5). Time traces undergoing single-step photobleaching events or displaying intensities from single molecules were selected for further analysis. After detecting the photobleaching events in iSMS, the photobleaching time and the fluorescence intensity of each trace were extracted using a custom MATLAB script.

**High-resolution optical tweezers experiments**

Nucleosome pulling experiments were carried out using a homebuilt dual trap optical tweezers instrument (6) by holding individual nucleoprotein complexes in between two polystyrene beads of 1 μm in diameter each. Trap stiffnesses were in the 0.4 – 0.7 pN/nm range. Raw data was acquired at 2.5 kHz. Purified single nucleosomes, hexasomes, and tetrasomes were ligated to 570 bp (5' end) and 700 bp (3' end) DNA handles at a concentration of 50 nM each, using *E. coli* DNA ligase (NEB) for 2h at 16˚C in the presence of 0.02% NP-40. The samples were subsequently ligated to carboxyl-functionalized polystyrene beads (Bangs labs), which were conjugated to double-stranded oligonucleotides (IDT) with one 5' NH-ester end (crosslinked to the bead) and one 4 nt sticky end (for handle ligation). The 4 nt sticky end was ligated with the 570 bp handle using *E. coli* DNA ligase at 16°C for 2 hr to attach nucleosomes and subnucleosomal particles to the bead. The free 700 bp handle was designed with a 5'-biotin to allow for later attachment to a streptavidin-coated polystyrene bead (Banglabs). Before the experiment, the tubing for nucleosome-bead injection was passivated for 30 min with 20 mM HEPES-KOH pH 7.5, 50 mM KOAc, 1 mg/mL BSA (NEB), followed by a second passivation for 30 min with 20 mM HEPES-KOH pH 7.5, 50 mM KOAc, 0.5% pluronic acid. Tethering was performed by trapping a single streptavidin bead near the nucleosome-ligated bead and allowing the biotin on the 700 bp handle to bind. Then, pulling and relaxing cycles were performed at 20 nm/s in pulling buffer with different $K^+$ concentrations (20 mM HEPES-KOH pH 7.0, 0/50/200 mM

KOAc, 5 mM $MgCl_2$, 0.1 mg/mL BSA, 0.02% NP-40, 10 mM $NaN_3$, 0.2 mM DTT). Sodium azide was used as oxygen scavenger system to increase tether lifetime.

**High-resolution optical tweezers with simultaneous FRET detection**

**Optical trapping instruments.** "Fleezers" experiments were performed in a custom-built instrument combining high-resolution optical traps and a single-molecule confocal fluorescence microscope modified from the design in Comstock *et al*. (7, 8). In this setup, a 1064 nm laser is passed through an acousto-optic deflector (AOD) with the laser alternating in position between the two traps every 5 µs. The detection of the bead positions in both traps was achieved on the same QPD. The position of the beads relative to the traps was measured using back focal plane interferometry (9). The confocal excitation laser (World Star Tech TECGL-30, 532 nm, 30 mW) was coupled through the right port of the microscope. The excitation laser was scanned by a piezo-controlled steering mirror (NanoDrive, MadCity Labs). The fluorescence emission was filtered from the infrared laser by a band pass filter and separated from excitation signals by a dichroic mirror, then was detected by two avalanche photodiodes (COUNT-10B APD for Cy3 fluorescence detection and COUNT-10 APD for Cy5 fluorescence, Laser Components). Data from the optical traps were collected at 1 kHz. Fluorescence data were initially recorded by the field-programmable gate array (FPGA) hardware with 1 ms integration, further integrated by software to 10 ms and saved. Three interlace cycles were used to generate two optical traps. To increase the observation window and avoid photobleaching, green laser and APD detection gate were interlaces and were turned on only in the interlace cycle when the trapping laser was off.

**Analysis of pulling and relaxation trajectories.** Each trajectory was properly corrected using the calibration factors obtained from the power spectrum fitting. For each trajectory, the force, extension, and fluorescence channels were recorded. Each trajectory was then segmented to separate the pulling trajectory (traps getting apart) from the relaxing trajectory (traps coming closer). Force and extension channels were used to plot force-extension curves, and the force at which a transition occurred as well as the corresponding change in extension were measured.

**FRET of unwrapping and relaxing trajectories.** The FRET efficiency ($E_{FRET}$) was calculated as the acceptor fluorescence intensity ($I_A$) divided by the sum of acceptor and donor fluorescence intensity ($I_D$) after background subtraction and cross-talk corrections for donor bleed-through to the acceptor channel.

$$E_{FRET} = \frac{I_A}{I_A + I_D}$$

**Alignment of fluorescent traces.** Alignment of FRET traces, where a clear force-transition was observed (high-force transition of ED2 and INT nucleosomes, and cooperative low-force transition of ED1 nucleosome), was performed by finding the rip location, which corresponded to the maximum change in force over time along the pulling curve, and setting the time of occurrence to zero. For the non-cooperative low-force transitions of ED1 nucleosome, a reverse logistic function fitting was performed for the FRET traces using the next equation:

$$f(x) = y_0 + \frac{a}{1 + e^{k(x-x_0)}}$$

where the fitting parameter $x_0$ was set to zero and subsequently used for alignment of the FRET traces.

**Pulling and relaxing experiments of fluorescently-labeled nucleosomes.** Fluorescently-labeled nucleosomes (H2A-Cy3 and Cy3/Cy5 nucleosomes) were ligated to 2.5 kb DNA handles upstream and downstream and attached to the oligo- and streptavidin-coated beads using the same strategy as in the high-resolution optical tweezers pulling experiments. Tubing for the nucleosome-bead was passivated as explained for high-resolution optical tweezers experiments. Pulling buffer consisted of 50 mM Tris-HCl pH 8.0, 50 mM KCl, 5 mM $MgCl_2$, 0.2 mg/mL BSA, 2% glucose, 0.02% NP-40, 2 mM TROLOX, 0.75 mg/mL Glucose Oxidase, 0.2 mg/mL Catalase (SIGMA), 0.4 mM DTT. Pulling buffer could be used for 60-90 min before its acidification. For a single experiment, a streptavidin bead was trapped in one trap and a nucleosome-DNA handle bead was trapped in the other, using low-intensity trapping lasers to increase fluorophore lifetime (~20% of laser power used to obtain a trap stiffness of 0.35 pN·nm). Once a tether was formed, laser power was increased to the value corresponding to ~0.35 pN·nm trap stiffness. Then, the tether was maintained under a tension of 1 pN, and a confocal scan using the green laser was performed to locate the position of the fluorescent sample. This position was later fixed for the rest of the samples analyzed. For pulling experiments, a symmetric pulling was performed at a speed of 50 or 100 nm/s until till force reached ~35 pN. The green laser was on during this whole period. Three pulling and relaxation cycles were performed per tethered molecule.

**SI Mechanical model of nucleosome unwrapping under force**

**Low-force transition.** The change in extension induced by nucleosome-rotation was measured using a simplified geometrical simulation. In detail, nucleosome crystal structure (PDB id: 1AOI; nucleosome wrapped by 146 bp of DNA) was rotated and transformed by aligning its dyad axis with the y-axis and the vector connecting the two DNA end points (weight center of DNA residue 1 and residue 146) with the x-axis using the VMD software (http://www.ks.uiuc.edu/Research/vmd/) (10). To incorporate the linker DNA into the model, an extra 40 bp of linear DNA was connected to each nucleosomal DNA end in the corresponding directions of extension (measured from a 10 bp distal segment on each side), forming the initial "crossing arms" conformation. The simulation of the low-force transition consisted of two stages; the simulation of rotation of a fully wrapped nucleosome, and the unwrapping of 27 bp at the weak DNA arm (distal in the 601 sequence) with simultaneous nucleosome rotation. During the first stage, the nucleosome model was rotated 144˚ in a total of 25 movie frames, while the extra 40 bp linkers were linearly interpolated to their target positions on the x-axis. At the second stage, single base pairs were sequentially removed from the crystal structure and added to the linker DNA which increases its length, while the torque generated at each round of distal DNA base-pair removal was eliminated, producing a residual rotation. This procedure was performed by repeating the following operation: i) calculation of the rotational matrix by aligning the vector that connects the entry and exit DNA points with the x-axis, ii) calculation of the translational matrix by aligning the center of the entry-exit points vector to the coordinate system origin, and iii) application of rotational and translation matrices to the whole nucleosome model. To keep track of DNA unwrapping during the whole simulation process, the two DNA ends were labeled with yellow marks, and the distance between them was measured using the VMD software. The 28 bp position on the weak DNA arm was labeled with red marker to mark the localization of the acceptor dye of the INT nucleosomal position.

**High-force transition.** The second transition of nucleosome unwrapping was performed using the procedure employed for the unwrapping at the low-force transition. The unwrapping at high-force begins with the model obtained at the end of the low-force transition (nucleosome wrapped by ~120 bp and with the torque of both DNA ends aligned to the x-axis). The unwrapping occurs symmetrically and alternately in 1 bp increments from both DNA ends until the more unwrapped weak DNA arm encounters the $(H3\text{-}H4)_2$ tetramer region, switching the unwrapping to only the strong DNA arm. Once the strong DNA arm reaches the tetramer region, the DNA unwrapping becomes symmetrical again and alternates between arms. This unwrapping process produces a nucleosome rotation of 180˚.

**SI Geometrical model of nucleosome unwrapping**

Force-extension curves obtained using high-resolution optical tweezers were down-sampled to 250 Hz for analysis. Rips and zips were analyzed using a custom MATLAB code. In short, a linear fitting of the data before and after the transition was performed. Later, a distance between the two linear equations $(\Delta x)$ is calculated from the point of the transition. We approximate the octamer core as a sphere of radius $r$. $\Delta x$ corresponds to the difference between the length of DNA interacting with the octamer core, $L_w$, minus the distance along the pulling axis between the last two points of contact of the DNA around the core (a secant) given by,

$$L_s = 2rsin\left(\pi - \frac{\theta}{2}\right) \qquad (1)$$

Assuming that the DNA wraps by an angle θ around de core, $L_w = \theta r$ and, therefore, $\Delta x$ upon the full straightening of the DNA under the applied force is,

$$\Delta x = L_w - L_s = r\left[\theta - 2sin\left(\pi - \frac{\theta}{2}\right)\right] \qquad (2)$$

By comparing the extension of the nucleosome pulling curve after the high-force transition and that of the bare DNA (Fig. S10A), we observe a difference $\Delta d$ of ~2.3 nm (s.d.= 0.6 nm). Using $r$ = 4.3 nm, (assuming an octamer core radius of 3.3 nm plus half of the DNA width of 1 nm), and applying Eq. (1) we find that $\alpha$ = 2.4 ± 0.2 radians (137 ± 11.5˚). Therefore, the length of DNA that remains interacting with the histone core after the high force transition is $L_{resid} = \alpha r$ = 10.3 ± 0.9 nm or 30 ± 3 bp.

The change in extension associated with the high-force transition (Fig. S10B), $\Delta x$ = 24.5 ± 1.6 nm corresponds to the difference between the sum of the length of left $(L_2)$ and right $(L_1)$ DNA arms unwrapped in the transition and the secant of the 30 ± 3 bp bound $(L_{s,resid})$, minus the distance along the pulling axis between the last two points of contact before the high force rip $(L_{s,before-HF})$, namely,

$$\Delta x = \left(L_2 + L_1 + L_{s,resid}\right) - L_{s,before-HF} \qquad (3)$$

The mechanical unwrapping model predicts that ~119 bp are bound before the high-force transition, which corresponds to the wrapping of DNA around the histone core by an angle of 9.4 radians. Therefore, after one full DNA turn (6.3 radians) is unwrapped, the remaining DNA forms an angle $\beta$ of 3.1 radians (177˚). Using, Equation (1) and $r$ = 4.3 nm, we find that $L_{s,before-HF}$ = 2*4.3**sin*(3.1/2) = 8.6 nm. Also, $L_{s,tet}$ = 2*4.3**sin*((2.4 ± 0.2)/2) = 8.0 ± 0.3 nm. Finally, using equation (3) and $\Delta x$ = 24.5 ± 1.6 nm, we find that,

$$(24.5 \pm 1.6 \text{ nm}) = L_1 + L_1 + (8.0 \pm 0.3) - (8.6 \text{ nm})$$

or, $L_{left-arm} + L_{rigth-arm}$ = 25.1 ± 1.6 nm. Indicating that during the high-force transition, around 25.1 ± 1.6 nm or ~74 ± 5 bp are unwrapped.

### Supporting Figures

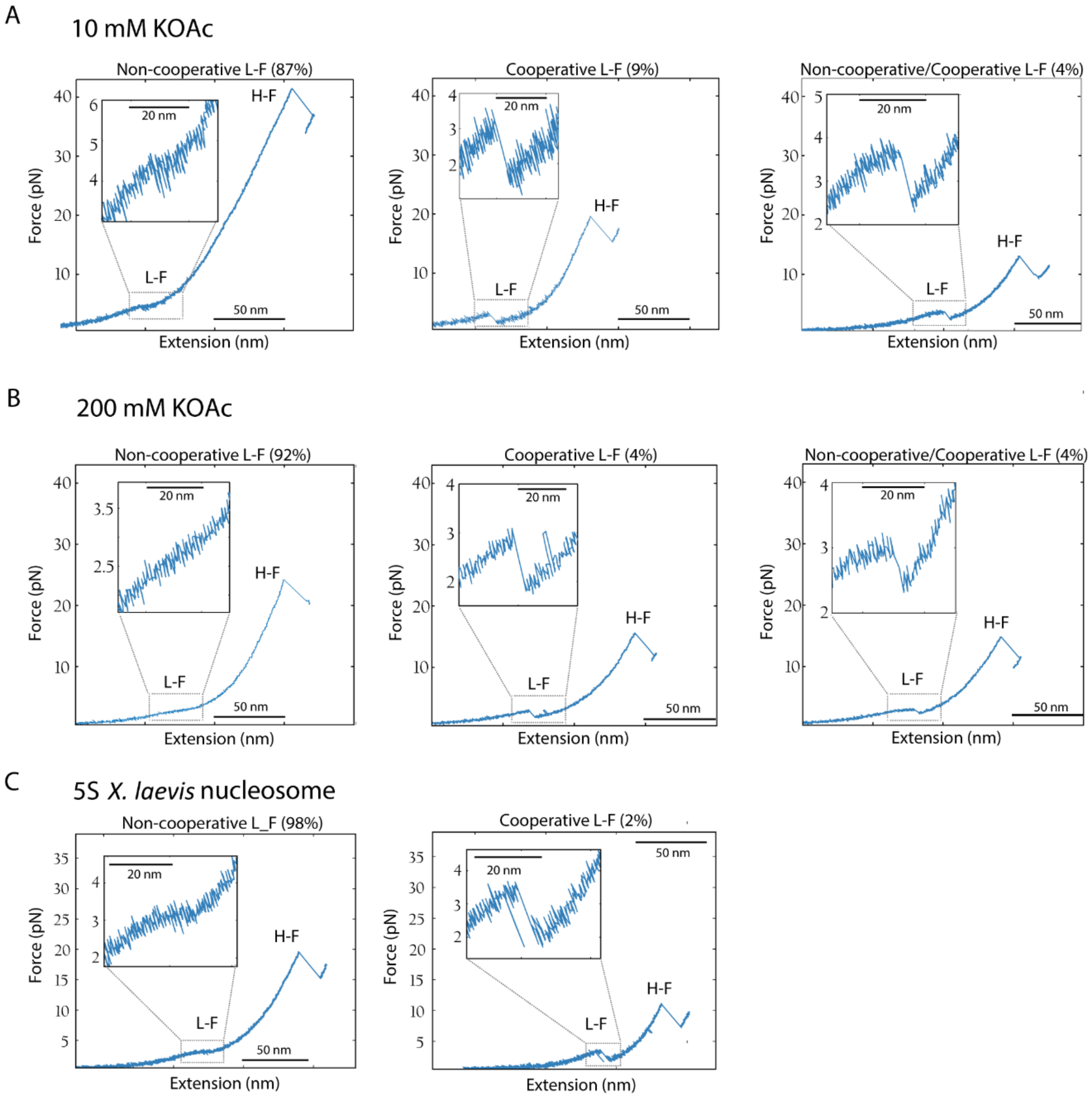


**Fig. S1. Force-extension unwrapping trajectories of *X. laevis* nucleosomes assembled on 601- and 5S-NPS.** First unwrapping trajectories of recombinant *X. laevis* nucleosomes showing the low-force transition (L-F) and the high-force transition (H-F). The low-force transition exhibits a non-cooperative, or cooperative, or both types of unwrapping trajectories. Percentage corresponds to the type of low-force trajectory. Insets correspond to a magnification of the low-force transition. (A) pulling curve of 601 nucleosome at 10 mM KOAc (from KOH). (B) Pulling curve of 601 nucleosome at 200 mM KOAc. In this high salt condition, low-force transition does not exhibit a cooperative trajectory. (C) Pulling curve of 5S nucleosome at 50 mM KOAc. 5S nucleosome does not exhibit a combination of both type of trajectories at low-force. Force and changes in extension values are summarize in Table S1 (low-force transition) and Table S2 (high-force transition).

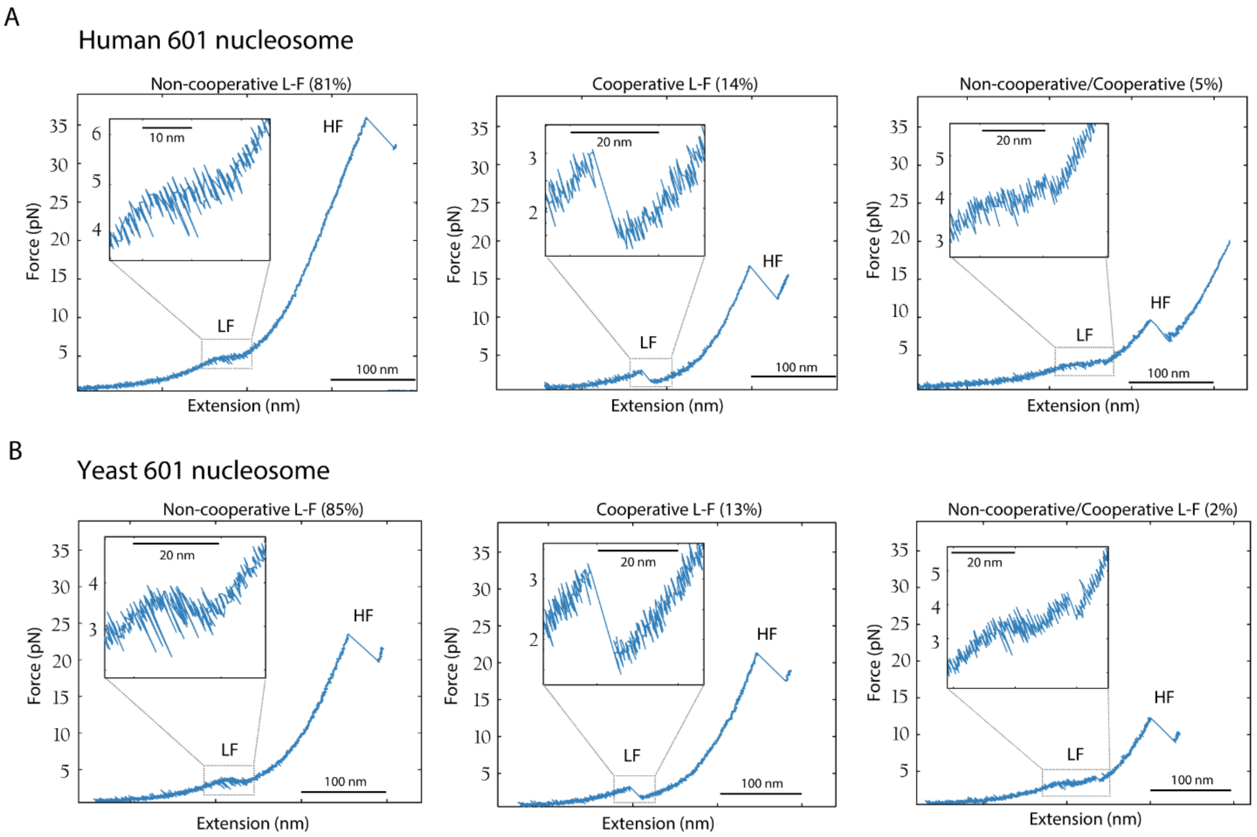


**Figure S2. Force-extension unwrapping trajectories of human and yeast 601 nucleosomes.** Pulling curves at 50 mM KOAc exhibiting the low-force transition (L-F) and the high-force transition (H-F). The low-force transition exhibits a non-cooperative, or cooperative, or both types of unwrapping trajectories. Percentage corresponds to the type of low-force trajectory. Insets correspond to a magnification of the low-force transition. (A) Recombinant human nucleosomes. (B) Recombinant yeast nucleosomes. Force and changes in extension values are summarize in Table S1 (low-force transition) and Table S2 (high-force transition).

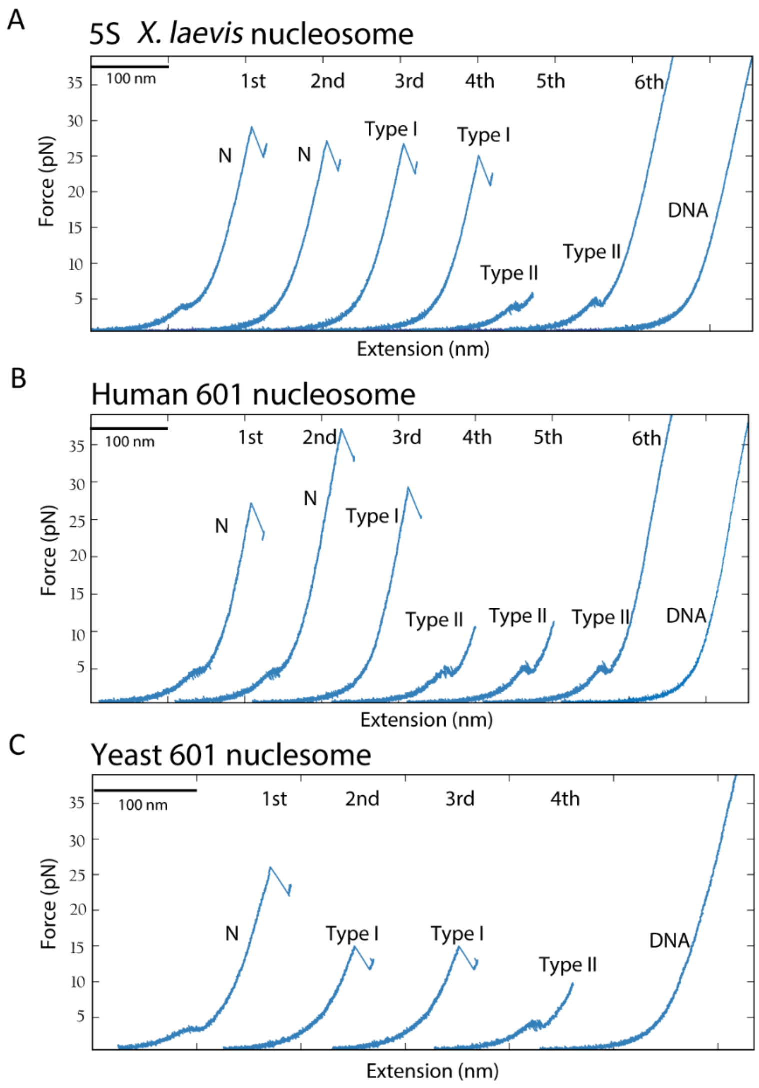


**Fig. S3. Mechanical disassembly of recombinant nucleosomes generates two intermediates.** Nucleosome disassembly by pulling and relaxation cycles (1st, 2nd, 3th…) at 50 mM KOAc of (A) *X. laevis* 5S nucleosome, (B) human 601 nucleosome, and (C) yeast 601 nucleosome. Only pulling curves (light blue) are shown and they were artificially separated for illustrative purposes. The first pulling curve from the first cycle correspond to the nucleosome (N). Type I intermediate exhibit a single rip at high force, and type II intermediate exhibit a single transition at a lower force. Bare DNA indicates full disassembly.

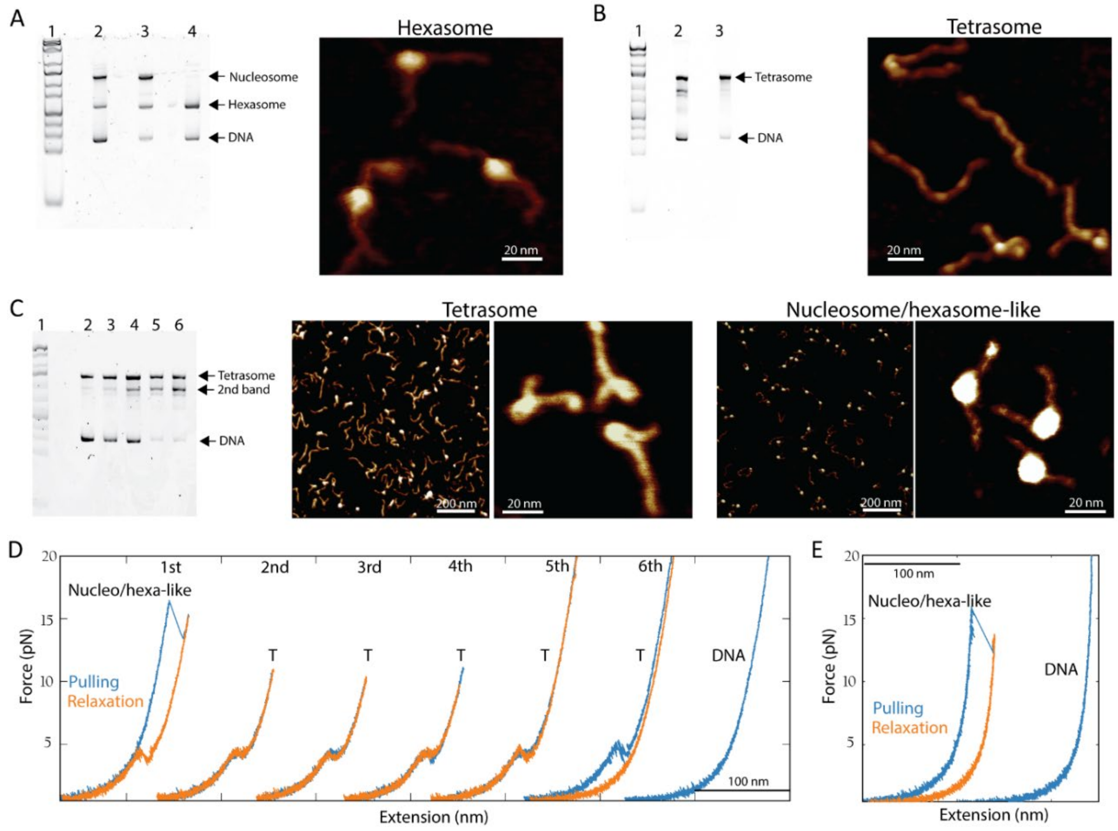


**Fig. S4. Atomic force microscopy imaging of hexasomes and tetrasomes.** (A) The assembly of *X. laevis* 601 nucleosomes by salt dialysis generates nucleosomes, hexasomes, and bare DNA (lane 2; left panel). Preparative purification using 5% acrylamide allowed to separate nucleosomes (lane 3) from hexasomes (lane 4) and reduce the amount of free DNA. Lane 1 corresponds to 1 kb plus DNA ladder. AFM image of pure hexasomes (right panel). In comparison to nucleosomes, hexasome DNA linkers extend from the core in the same plane and in opposite directions and they exhibit a similar length. Hexasome average maximum height correspond to ~3 nm (n= 4). (B) *X. laevis* 601 tetrasomes were assembled using pure $(H3\text{-}H4)_2$ tetramers in excess of 1.4-fold compared to DNA (lane 2), and the resulting product was purified by preparative electrophoresis (lane 3). Lane 1 corresponds to 1 kb plus DNA ladder. AFM imaging of pure tetrasomes shows that tetrasome keeps the curvature on 601 NPS, and exhibits an average maximum height of ~1.5 nm (n= 3). (C) H3-H4 tetramer assembly at increasing concentration of H3-H4 relative to DNA, produces two main bands as visualized by 5% native electrophoresis (left panel. Lane 1= 1 kb plus ladder; lane 2= 1:1; lane 3= 1:1.4; lane 4= 1:1.8; lane 5= 1:2.2; lane 6= 1:2.6. Ratios correspond to DNA:$(H3\text{-}H4)_2$). The top band correspond to the tetrasome previously purified in (B). The bottom band (2nd band) increases its intensity at higher $(H3\text{-}H4)_2$ to DNA ratios. At a ratio of 1:1.4 DNA:$(H3\text{-}H4)_2$ (middle panel), pure tetramers form mainly tetrasomes and also few rounded and bigger particles resembling nucleosomes and/or hexasomes. At 1:2.6 DNA:$(H3\text{-}H4)_2$ ratio, the formation of the nucleosome/hexasome-like particles is favored (right panel), and they exhibit a height of ~3.5 nm (n= 3). (D) Unwrapping trajectories of nucleosome/hexasome-like particles exhibit a single cooperative force-extension transition centered at ~17 pN, which can disassembly into tetrasomes (T) by pulling (light blue curves) and relaxation (orange curves) cycles (1st to 6th) previous to full disassembly into DNA, or (E) directly into DNA after just one cycle of pulling and relaxation.

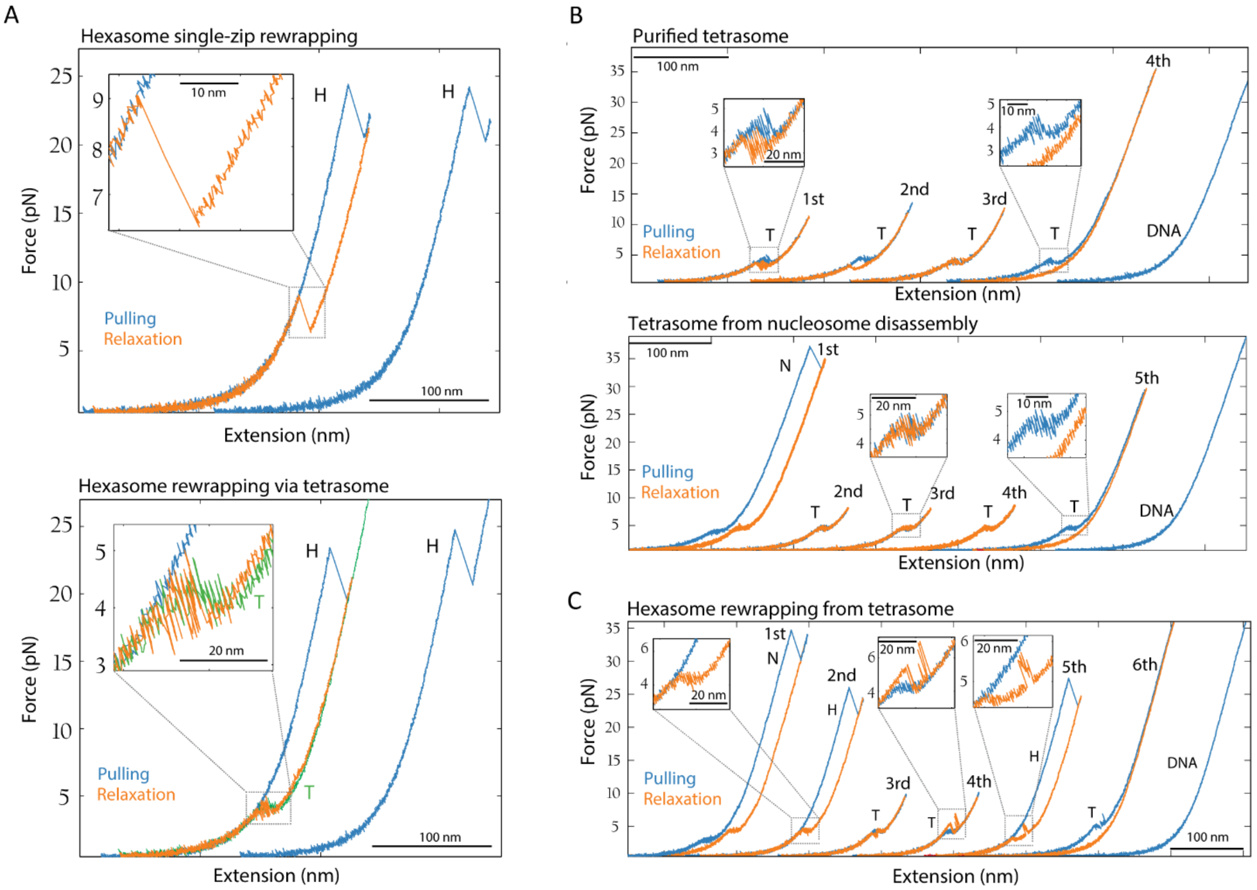


**Fig. S5. Reversible assembly of hexasomes and tetrasomes.** (A) Rewrapping of hexasomes (H) was characterized by analyzing the relaxation trajectory (orange curve) obtained between two hexasome pulling curves (light blue curves). The second pulling curve has been displaced laterally for clarity. Two main types of rewrapping trajectories were observed. The first type exhibits a single zip at ~6 pN (top panel). This single zip reaches the hexasome pulling curve and it has a change in extension of ~22 nm, similar to the extension observed for the hexasome high-force rip, and indicates that the hexasome rewrapping can occur in a single cooperative step. A second type of rewrapping curve shows first a continuous shortening that appears as a deviation from the worm-like chain behavior followed by fluctuations or hoping and a single zip at ~4 pN, which reaches the hexasome pulling curve (bottom panel). The zip size is ~17 nm; the shortening that precedes it corresponds to non-cooperative rewrapping of ~5 nm. This type of hexasome rewrapping trajectory passes through a tetrasome rewrapping intermediate (T; green curve). (B) The unwrapping and rewrapping of pure tetrasomes (top panel) or tetrasomes derived from nucleosome (N) disassembly (bottom panel) is reversible, but in some cases, they exhibit different degrees of force-extension hopping (insets). Cycles of pulling and relaxation (1st, 2nd, 3th, $4^{th}$, $5^{th}$) were artificially separated for illustrative purposes. (C) In few cases, tetrasomes that have been generated by nucleosome disassembly in which the pulling and relaxation cycles did not exceed ~10 pN, it was observed pulling curves going back ($5^{th}$ cycle) to those characteristics of the unrewrapping of hexasomes, displaying a single rip at high force. These observations suggest that H2A-H2B heterodimers can remain bound to the DNA during unwrapping and eventually re-engage the tetrasome.

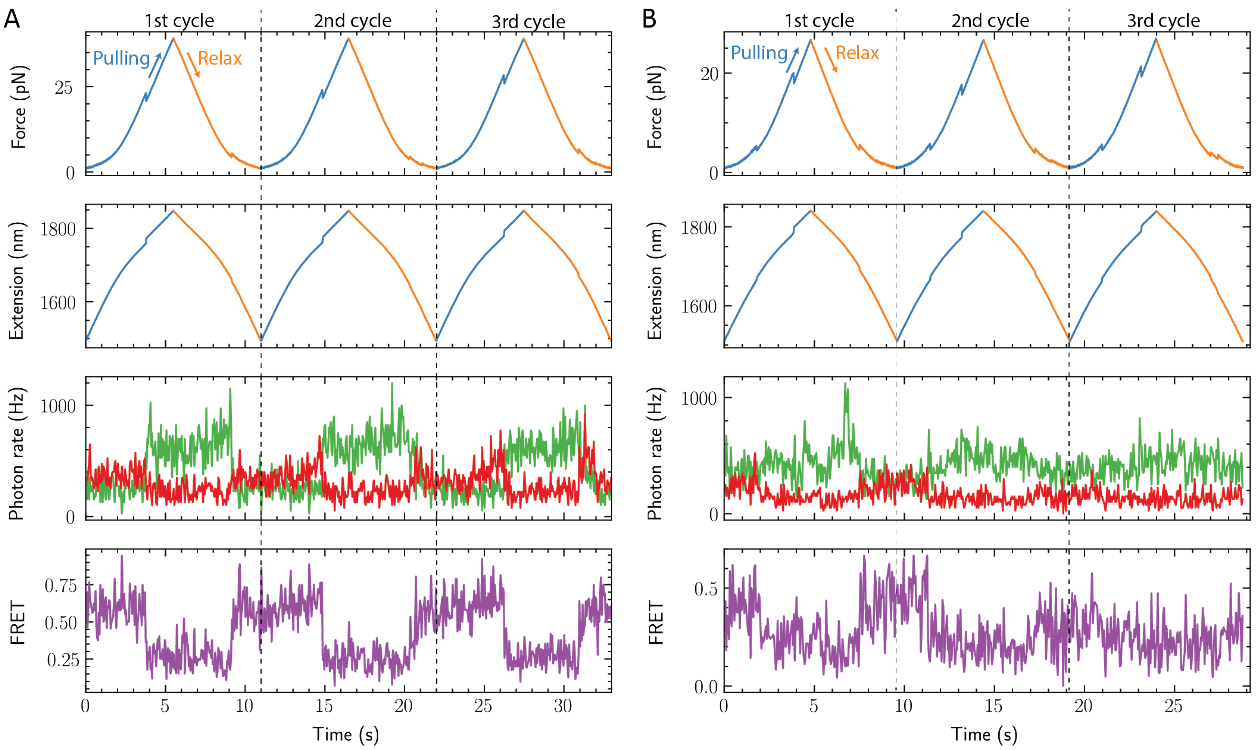


**Fig. S6. Mechanical unwrapping of INT and ED2 nucleosomes monitored by co-temporal force and fluorescence measurement.** Simultaneous force, extension, and fluorescence measurements of (A) INT, and (B) ED1 nucleosomes during three pulling and relaxation cycles (separated by black dashed lines) under 532 nm green laser excitation. Blue and orange traces indicate puling and relaxation cycles, respectively. Fluorescence channel detects the anticorrelated changes in green and red signals corresponding to changes in FRET. Distinctive and simultaneous transitions in the force, extension, and FRET occur at high-force (~20 pN) and low-force (~5 pN) for INT and E1 nucleosomes, respectively. The recovery of the FRET signal indicates nucleosome rewrapping.

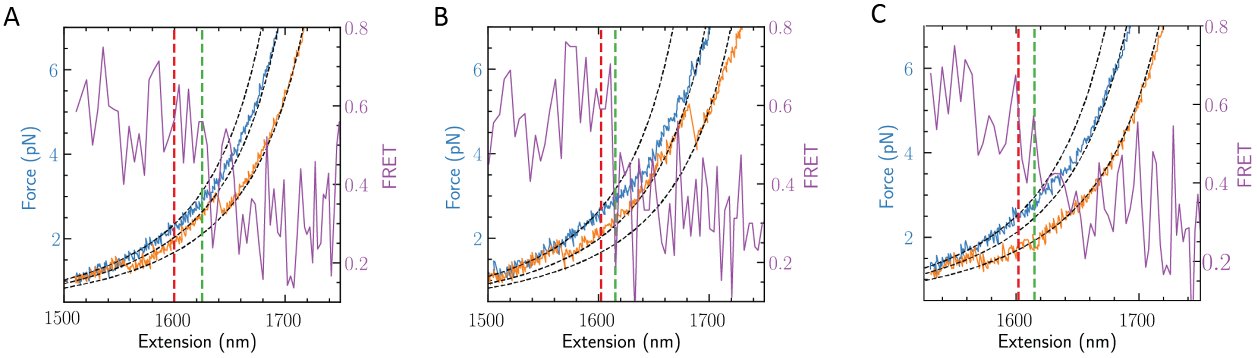


**Fig. S7. Non-cooperative low-force transition and FRET decrease are not synchronized for ED1 nucleosome.** (A-C) Three examples of ED1 nucleosomes showing the temporal relation between the non-cooperative decrease in FRET as a function of distance (purple trace) and the non-cooperative change in extension of ED1 nucleosomes. The fitting of the worm-like-chain model (dashed black lines) to the low-force section of the nucleosome pulling curve (blue solid line) exhibit a divergence as a consequence of the beginning of the non-cooperative low-force transition (red dashed line). Hexasome relaxation trajectory (orange curve) was included as a reference to validate the worm-like-chain fitting.

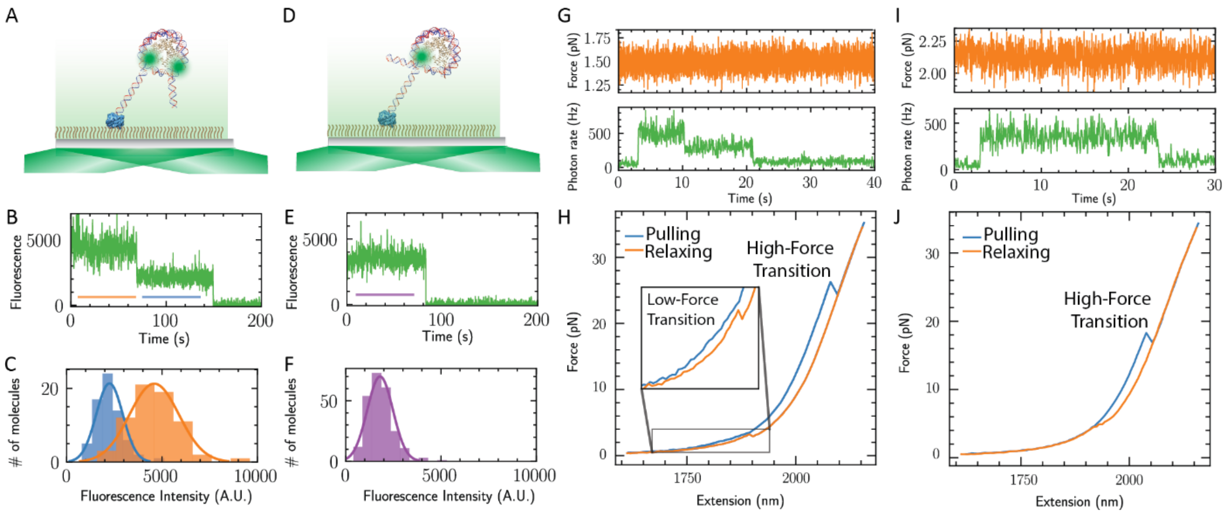


**Fig. S8. Single-molecule fluorescence characterization of the nucleosome's integrity and histone stoichiometry.** (A) Schematics of smTIRF experiments of H2A-Cy3 nucleosomes. Nucleosomes were attached to the streptavidin coated surface using a biotin molecule at one DNA end. (B) Time course of a single fluorescent nucleosome showing a two-step photobleaching indicate that they harbor two H2A-Cy3. (C) Fluorescence intensity distribution corresponding to the first step bleaching (orange) and second step bleaching (blue). (D) Schematics of smTIRF experiments of H2A-Cy3 hexasomes. (E) Time course of a single fluorescent hexasome showing a single-step photobleaching confirms the presence of a single H2A-Cy3. (F) Fluorescence intensity distribution corresponding to the single step bleaching (purple). Note that the fluorescence intensity of H2A-Cy3 nucleosomes (orange in Fig. S8C) is twice that of H2A-Cy3 hexasomes whose signals are comparable to those of nucleosomes after one photobleaching event (blue in Fig. S8C). (G) Co-temporal force and fluorescence measurements of a single tethered H2A-Cy3 nucleosome. The tether was held at ~1.5 pN where no mechanical transition on the nucleosome is observed. Two photobleaching steps indicate the presence of two H2A-H2B heterodimers in the nucleosome. (H) Force-extension unwrapping trajectory of H2A-Cy3 nucleosome exhibit the low- and high-force transitions. (I) Co-temporal force, extension, and fluorescence measurements of a single tethered H2A-Cy3 hexasome. A single photobleaching step indicate the presence of one H2A-H2B heterodimer in the hexasome. (J) Force extension unwrapping trajectory of H2A-Cy3 hexasome exhibit the single rip transition.

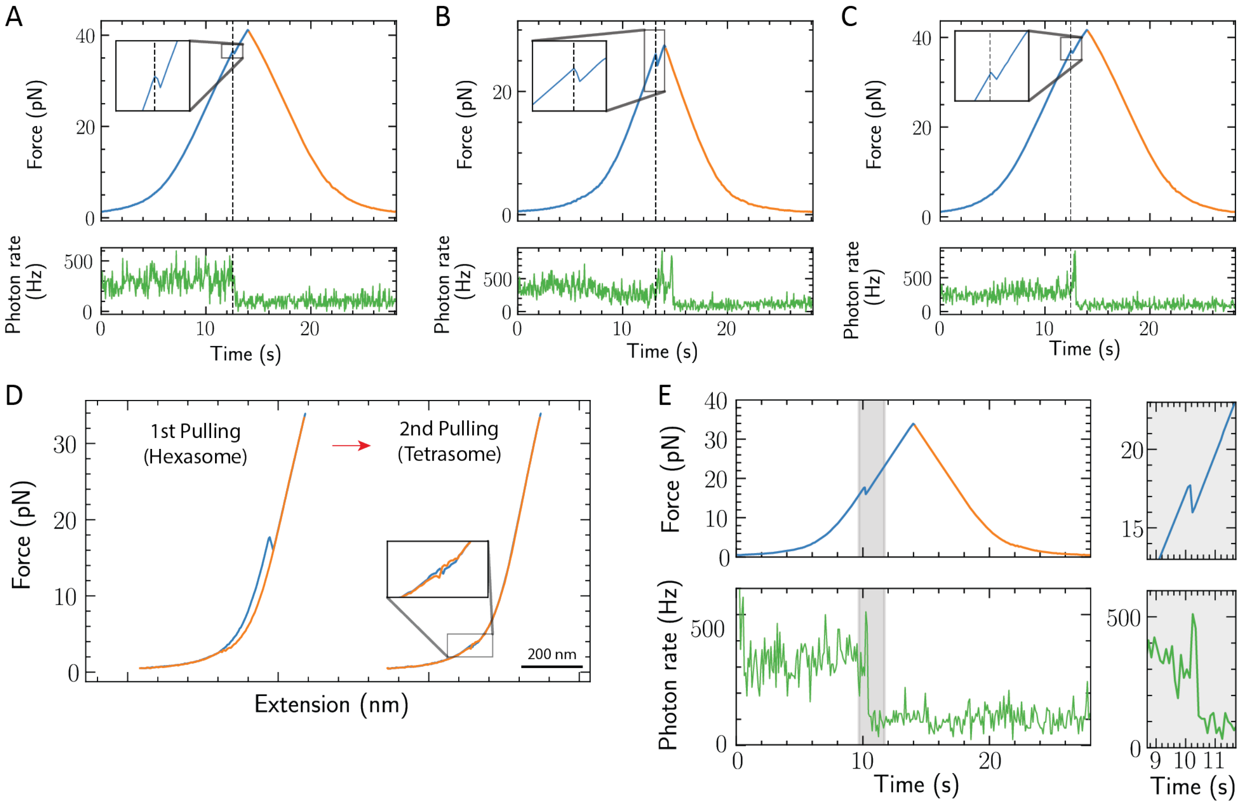


**Fig. S9. Real-time detection of H2A-H2B heterodimer dissociation.** (A-C) Simultaneous time course of the fluorescence and force channels during the first pulling (blue) and relaxation (orange) of three different H2A-Cy3 nucleosomes. Inset correspond to a magnification of the high-force extension transition. (D) H2A-Cy3 hexasome disassembly by two cycles of pulling (blue curve) and relaxation (orange curve). Pulling cycles were artificially separated for illustrative purposes. The single low-force transition in the second pulling curve indicates the formation of a tetrasome after hexasome unwrapping. (E) Simultaneous time course of the fluorescence and force channels during the first unwrapping curve of the hexasome in (D). Shaded gray area indicates the zoom regions of the plots shown to the right.

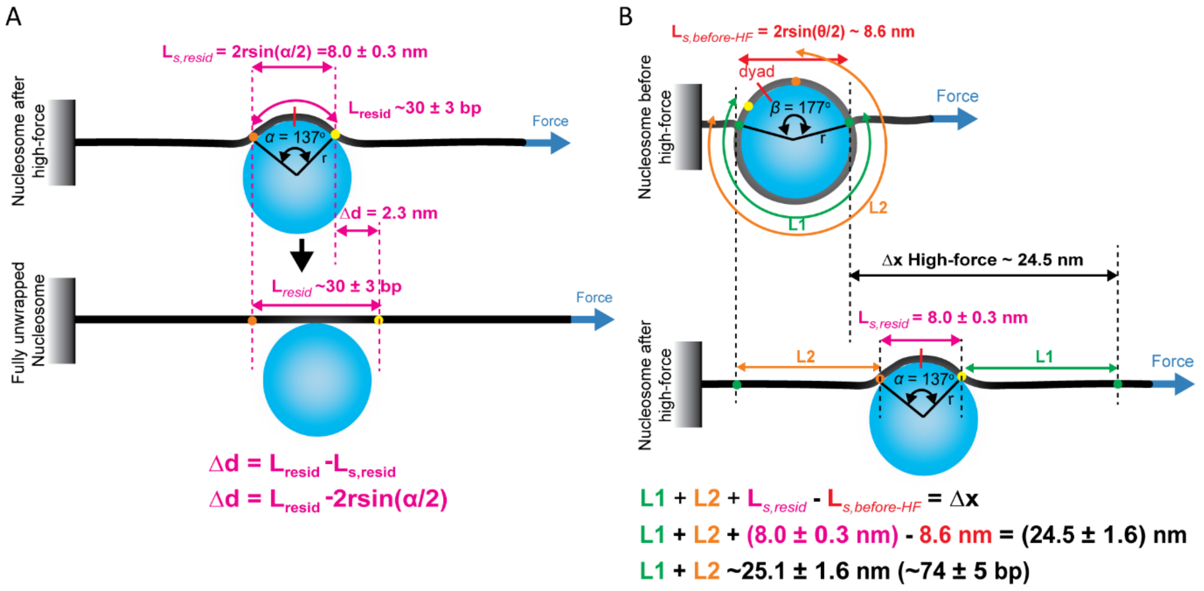


**Figure S10. Geometrical model of nucleosome unwrapping.** For details of see SI Materials and Methods. DNA is depicted in black, while the histone core is represented as a sphere (cyan). (A) Upon the application of force (blue arrow), the residual DNA bound to the histone core ($L_{resid}$ = 30 ± 3 bp, purple) unwraps, producing a change in extension of $\Delta d$ = 2.3 ± 0.6 nm. The distance between the last two point of contacts (orange and yellow dots) corresponds to $L_{s,resid}$ = 8.0 ± 0.3 nm (secant). (B) Unwrapping at the high-force transition ($\Delta x$). The distance between the last two points of contacts (green dots) of the DNA bound to the core is equivalent to the *secant* $L_{s,before\text{-}HF}$ ~8.6 nm (top panel). Unwrapping of the left and right DNA arms corresponds to L2 (orange curve, between the green and orange dots) and L1 (green curve, between green and yellow dots), respectively (bottom panel). At the end of the high-force transition, the secant of the residual DNA bound to the histone core corresponds to $L_{s,resid}$ = 8.0 ± 0.3 nm.

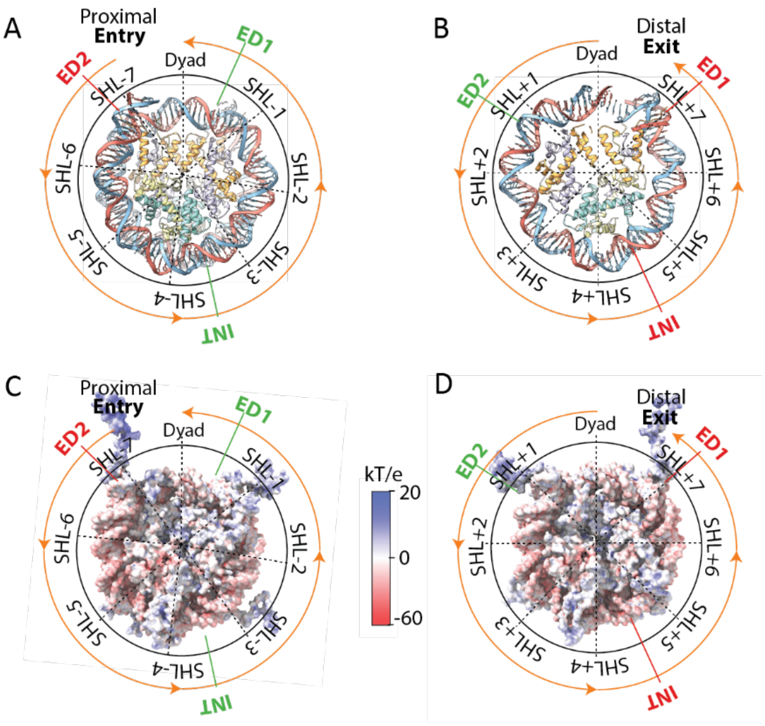


**Fig. S11. Overview of nucleosome structure and its electrostatic surface potential.** Nucleosome structure displaying Super-Helical Locations (SHLs) from the (A) proximal and (B) distal side (PDB: 6ESF). Red and Green indicators on the periphery indicate the location of the donor (green) and acceptor (red) of the FRET pairs of ED2, ED1, and INT nucleosomes. Poisson-Boltzmann calculations of the nucleosome electrostatic surface potential (PDN: 1KX5) with views from the (C) proximal and (D) distal side. The electrostatic potential mapped to the molecular surface was calculated using Adaptive Poisson-Boltzmann Solver (APBS) (11). The electrostatic potential ranges from +20 (blue) to −60 (red) kTe−1.

**Table S1. Force-extension parameters of the low-force transition of recombinant nucleosomes**

| | | | Non-cooperative | | Cooperative | | | Non-cooperative/cooperative | | |
|---|---|---|---|---|---|---|---|---|---|---|
| **Nucleosome** | **KOAc (mM)** | **N** | **Force (pN)** | ***n* (%)** | **Force (pN)** | **Change in ext. (nm)** | ***n* (%)** | **Force (pN)** | **Change in ext. (nm)** | ***n* (%)** |
| ***X. Laevis* 601 NPS** | 10* | 54 | 4.3 ± 0.3 | 47 (87) | 3.4 ± 0.5 | 23.5 ± 1.6 | 5 (9) | 4.2 ± 0.2 | 18.8 ± 4.9 | 2 (4) |
| ***X. Laevis* 601 NPS** | 50 | 53 | 3.9 ± 0.3 | 46 (87) | 4.1 ± 0.7 | 20.4 ± 1.5 | 5 (9) | 3.8 ± 0.2 | 20.6 ± 2.0 | 2 (4) |
| ***X. Laevis* 601 NPS** | 200 | 50 | 3.2 ± 0.4 | 46 (92) | 2.6 ± 0.3 | 20.8 ± 0.9 | 2 (4) | 3.0 ± 0.1 | 26.7 ± 2.4 | 2 (4) |
| ***X. Laevis* 5S-NPS** | 50 | 36 | 3.8 ± 0.3 | 35 (98) | 3.4 | 22 | 1 (2) | ND | ND | ND |
| **Human 601 NPS** | 50 | 43 | 4.5 ± 0.3 | 35 (81) | 3.9 ± 0.5 | 21.9 ± 1.4 | 6 (14) | 4.8 ± 0.2 | 17.6 ± 0.1 | 2 (5) |
| **Yeast 601 NPS** | 50 | 61 | 3.5 ± 0.3 | 52 (85) | 3.5 ± 0.3 | 22.5 ± 1.9 | 8 (13) | 3.1 | 22 | 1 (2) |

ND= non-determined; NPS= nucleosome positioning sequence. *10 mM comes from the KOH used to titrate the buffer, no KOAc was added. Uncertainties are standard errors of the means.

**Table S2. Force-extension parameters of the high-force transition of recombinant nucleosomes**

| | **N** | **KOAc (mM)** | **Force (pN)** | **Change in ext. (nm)** |
|---|---|---|---|---|
| ***X. Laevis* (601 NPS)** | 56 | 10* | 34.8 ± 9.2 | 24.5 ± 1.1 |
| ***X. Laevis* (601 NPS)** | 54 | 50 | 30.4 ± 8.3 | 24.5 ± 1.6 |
| ***X. Laevis* (601 NPS)** | 48 | 200 | 24.4 ± 4.2 | 24.5 ± 1.2 |
| ***X. Laevis* (5S NPS)** | 32 | 50 | 29.4 ± 6.7 | 24.1 ± 1.6 |
| **Human (601 NPS)** | 43 | 50 | 28.9 ± 7.9 | 24.6 ± 1.7 |
| **Yeast (601 NPS)** | 62 | 50 | 21.1 ± 6.8 | 24.0 ± 1.1 |

NPS= nucleosome positioning sequence. *This pulling buffer did not contain potassium acetate. 10 mM comes from the KOH used to titrate the buffer. Uncertainties are standard errors of the means.

**Table S3. Nucleosome disassembly probabilities at different ionic forces**

| KOAc (mM) | | | | | | |
|---|---|---|---|---|---|---|
| | **0** | | **50** | | **200** | |
| **Second pulling** | **Probability** | ***n*** | **Probability** | ***n*** | **Probability** | ***n*** |
| Nucleosome | 0.283 ± 0.051 | 15 | 0.412 ± 0.062 | 21 | 0.286 ± 0.054 | 14 |
| Hexasome | 0.340 ± 0.051 | 18 | 0.294 ± 0.056 | 15 | 0.388 ± 0.060 | 19 |
| Tetrasome | 0.208 ± 0.043 | 11 | 0.275 ± 0.062 | 10 | 0.286 ± 0.054 | 14 |
| DNA | 0.169 ± .048 | 9 | 0.019 ± 0.104 | 1 | 0.040 ± 0.098 | 2 |
| | | 53 | | 47 | | 49 |

After the first pulling and relaxation cycle of *X. laevis 601* nucleosomes, the second pulling curve was classified as nucleosome rewrapping (two force transitions), hexasomes (single rip), tetrasomes (single low-force transition), and bare DNA. Probability of conversion was determined using the Bootstrap method.
